# Changes in Cell Size and Shape During 50,000 Generations of Experimental Evolution with *Escherichia coli*

**DOI:** 10.1101/2020.08.13.250415

**Authors:** Nkrumah A. Grant, Ali Abdel Magid, Joshua Franklin, Yann Dufour, Richard E. Lenski

**Affiliations:** Department of Microbiology and Molecular Genetics, Michigan State University, East Lansing, MI 48824; BEACON Center for the Study of Evolution in Action, Michigan State University, East Lansing, MI 48824; Department of Ecology, Evolutionary Biology, and Behavior Program, Michigan State University, East Lansing, MI 48824

## Abstract

Bacteria adopt a wide variety of sizes and shapes, with many species exhibiting stereotypical morphologies. How morphology changes, and over what timescales, is less clear. Previous work examining cell morphology in an experiment with *Escherichia coli* showed that populations evolved larger cells and, in some cases, cells that were less rod-like. That experiment has now run for over two more decades. Meanwhile, genome sequence data are available for these populations, and new computational methods enable high-throughput microscopic analyses. Here, we measured stationary-phase cell volumes for the ancestor and 12 populations at 2,000, 10,000, and 50,000 generations, including measurements during exponential growth at the last timepoint. We measured the distribution of cell volumes for each sample using a Coulter counter and microscopy, the latter of which also provided data on cell shape. Our data confirm the trend toward larger cells, while also revealing substantial variation in size and shape across replicate populations. Most populations first evolved wider cells, but later reverted to the ancestral length-to-width ratio. All but one population evolved mutations in rod-shape maintenance genes. We also observed many ghost-like cells in the only population that evolved the novel ability to grow on citrate, supporting the hypothesis that this lineage struggles with maintaining balanced growth. Lastly, we show that cell size and fitness remain correlated across 50,000 generations. Our results suggest larger cells are beneficial in the experimental environment, while the reversion toward ancestral length-to-width ratios suggests partial compensation for the less favorable surface area-to-volume ratios of the evolved cells.

**Importance:** Bacteria exhibit great morphological diversity, yet we have only a limited understanding of how their cell sizes and shapes evolve, and of how these features affect organismal fitness. This knowledge gap reflects, in part, the paucity of the fossil record for bacteria. Here, we revive and analyze samples extending over 50,000 generations from 12 populations of experimentally evolving *Escherichia coli* to investigate the relation between cell size, shape, and fitness. Using this “frozen fossil record” we show that all 12 populations evolved larger cells concomitant with increased fitness, with substantial heterogeneity in cell size and shape across the replicate lines. Our work demonstrates that cell morphology can readily evolve and diversify, even among populations living in identical environments.

## Introduction

For well over 100 years, cell biologists have wondered why cells adopt characteristic shapes (1) and sizes (2). Cell size has been of particular interest owing to its importance for organismal fitness. For example, cell size influences a bacterium’s susceptibility to predation by protists (3, 4) and phagocytosis by host immune cells (5, 6). Larger cell size has also been implicated in increasing susceptibility to bacteriophages (7, 8) and reducing susceptibility to antibiotics (9, 10, 11). Cell size is generally tightly coupled to growth and division. Most eukaryotic cells follow a four-stage cycle in which they must reach a critical mass before partitioning into daughter cells (12). In contrast, the bacterial cell cycle involves less discrete periods due to the overlapping nature of cell growth, DNA replication, chromosome segregation and division (13). Bacterial cells are generally larger when they are growing faster (14, 15, 16), in order to accommodate more genetic material (17, 13) and ribosomes (18). These facts suggest that cell size per se is a direct target of selection. However, it has also been suggested that cell size is a “spandrel” (19), i.e., a phenotypic character that might appear to be the product of adaptive evolution but is instead merely a byproduct of natural selection acting on some other trait (20).

The distribution of cell size in prokaryotes spans many orders of magnitude (21). The smallest known bacteria occur in the genus *Palagibacterales*; they constitute 25% of all marine planktonic cells, and they have average volumes of only ∼0.01 fL (1 fL = 1 µm^3^) (22, 23). The largest heterotrophic bacteria, in the genus *Epulopiscium*, live in the intestines of surgeonfish; they have cytoplasmic volumes of ∼2 x 10^6^ fL (24, 25). In contrast to these extremes, the average cell volumes of four widely studied bacteria— *Bacillus subtilis, Staphylococcus aureus, Escherichia coli, and Caulobacter crescentus—*range between about 0.4 – 3.0 fL (25).

Large bacterial cells face significant challenges. Unlike multicellular eukaryotes that use elaborate vasculature or similar systems to transport nutrients and waste between cells, along with specialized cells to acquire nutrients from and dispose of wastes to the environment, bacteria rely on diffusion to grow and reproduce (26, 27, 28). Diffusion must be considered from two perspectives. A cell must first acquire nutrients from the environment at the cell surface, and those nutrients must then diffuse internally to their sites of biochemical processing in a timely fashion. As cells grow, volume generally increases faster than surface area, such that the surface area-to-volume (SA/V) ratio decreases. The SA/V ratio might thus constrain viable cell sizes, as cells that are too large may be unable to obtain nutrients at a sufficient rate to service the demands of their biomass. However, bacteria have evolved a number of strategies that increase their rates of nutrient acquisition for a given cell volume. Rod-shaped cells, for example, experience a smaller reduction in their SA/V ratio as they grow larger than do spherical cells. Other examples include various extracellular projections that allow surface attachment while generating biomechanical motion to refresh the medium in the cell’s immediate environment (29); chemotaxis, which allows cells to move along gradients of increasing nutrient concentration (30); and invaginated cell envelopes, which increase the SA/V ratio (31).

The surface area of a spherocylindrical (i.e., rod-shaped) cell is given by 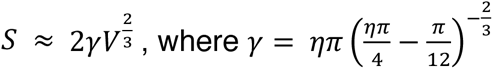 and η is the aspect ratio, i.e., the cell’s length divided by its width (32). For rod-shaped species like *E. coli*, the SA/V ratio can be varied by changing either a cell’s length or width, while holding the volume constant. Assuming that the rod shape is maintained, doubling a cell’s width reduces its SA/V ratio by much more than doubling its length (33). If all else were equal, then SA/V considerations would predict relatively larger cells during nutritional upshifts (resources plentiful) and smaller cells during nutritional downshifts (resources scarce).

Now, suppose a bacterial population resides in a simple environment, one free of predators and stressors and with a predictable supply of carbon. As this population adapts to this environment by natural selection, the cells grow slightly faster. Do the cells also become larger, and if so, how much larger? Does cell size constrain the maximum growth rate that can be achieved, or is the causality in the opposite direction? If the cells evolve to become larger, are they larger while growing, in stationary phase when the limiting resource is depleted, or both? And what might change about the shapes of the cells including their aspect and SA/V ratios?

Experimental evolution has proven to be powerful for addressing such questions. This research framework provides the opportunity to study evolution in real time, both in biological (34, 35, 36) and digital (37, 38) systems. In the long-term evolution experiment (LTEE), 12 replicate populations of *E. coli* were started from a common ancestor and have been propagated by daily serial transfer in a minimal glucose-limited medium for more than 70,000 generations (32 years). Whole-population samples, and clones from each population, have been frozen every 500 generations, creating a frozen “fossil record” from which genotypic and phenotypic changes can be measured (39, 40).

Evolution proceeded most rapidly early in the LTEE. By 2,000 generations, the populations were, on average, ∼35% more fit than their ancestor. An increase in the exponential growth rate and a reduction in the duration of the lag phase prior to growth were major contributors to this improvement (41). By 50,000 generations, the average population was ∼70% more fit than the ancestor (42, 43, 44). The trajectory for fitness relative to the ancestor is well described a power-law function, which implies that fitness may continue to increase indefinitely, albeit at progressively slower rates of improvement (42).

In the first 10,000 generations, cell size was found to have increased in all 12 LTEE populations and their trajectories were positively correlated with fitness (40). The increase in cell volume was accompanied by a concomitant decrease in numerical yield, although the product of cell volume and number—the total biovolume yield—increased (41). In the meantime, several populations were found to have diverged in shape, producing more spherical cells (45, 46) and fitness has continued to increase for at least 50,000 generations more (42, 43).

In this study, we sought to determine if cell size has continued to increase over time, and whether it still tracked with fitness. To that end, we measured cell size in the ancestor and the evolving populations at 2,000, 10,000 and 50,000 generations. We used both a Coulter particle counter and microscopy to measure cell volumes, and microscopy to characterize changes in cell shape. All 12 populations evolved larger cells. As previously seen in the fitness trajectories, the rate of change in cell volumes was fastest early in the experiment, and the trend was monotonically increasing over time. By 50,000 generations, the average cell volume in most populations was well over twice that of the ancestor, both during exponential growth and in stationary phase. The evolved cells tended to increase more in width than in length during the first 10,000 generations, but they subsequently reverted to aspect ratios similar to the ancestral strain. However, there was considerable among-population variability in shape as well as size through the entire period. Analyses of genome sequence data also revealed mutations in cell-rod maintenance genes in almost every population. Lastly, we discovered greatly elevated cell mortality in the only population that evolved the novel ability to use citrate in the growth medium as a carbon source. Overall, our data suggest that cell size and shape are important targets of selection in the LTEE.

## Results

Our analyses and results are multi-faceted. They include: a comparison between two methods used to measure cell volumes; analyses of the evolutionary trends in cell size of both clones and whole-population samples; a comparison of sizes during exponential growth and at stationary phase; analyses of cell shape and the subsequent identification of mutations in genes known to affect cell shape; the correlation between cell size and relative fitness during the LTEE; and evidence for substantial cell mortality in a unique population.

### Cell volumes measured by two methods

We first address whether the two approaches we used to estimate cell size provide comparable results. The Coulter-counter method directly estimates particle volumes, based on changes in conductance between two electrodes as cells suspended in an electrolyte solution are moved through a small aperture. The microscopy method involves obtaining cell images and processing them using software that defines the edges of objects, segments the objects into small pieces, and integrates the segments to estimate cell volumes. Fig. 1 shows the highly significant correlation in the median cell volumes estimated using the two approaches for the two ancestors and 36 evolved clones from the 12 populations at three generations of the LTEE. All of these samples were grown in the same LTEE conditions and measured in stationary phase at the 24-h mark (i.e., when they would be transferred to fresh medium under the LTEE protocol). This concordance gives us confidence that we can use either approach when it is best suited to a given question. The Coulter counter method is especially well suited to efficient measurement of cell volumes for many cells from each of many samples. The microscopy and subsequent image processing, by contrast, is necessary to obtain information on changes in cell shape.

**FIG 1.**
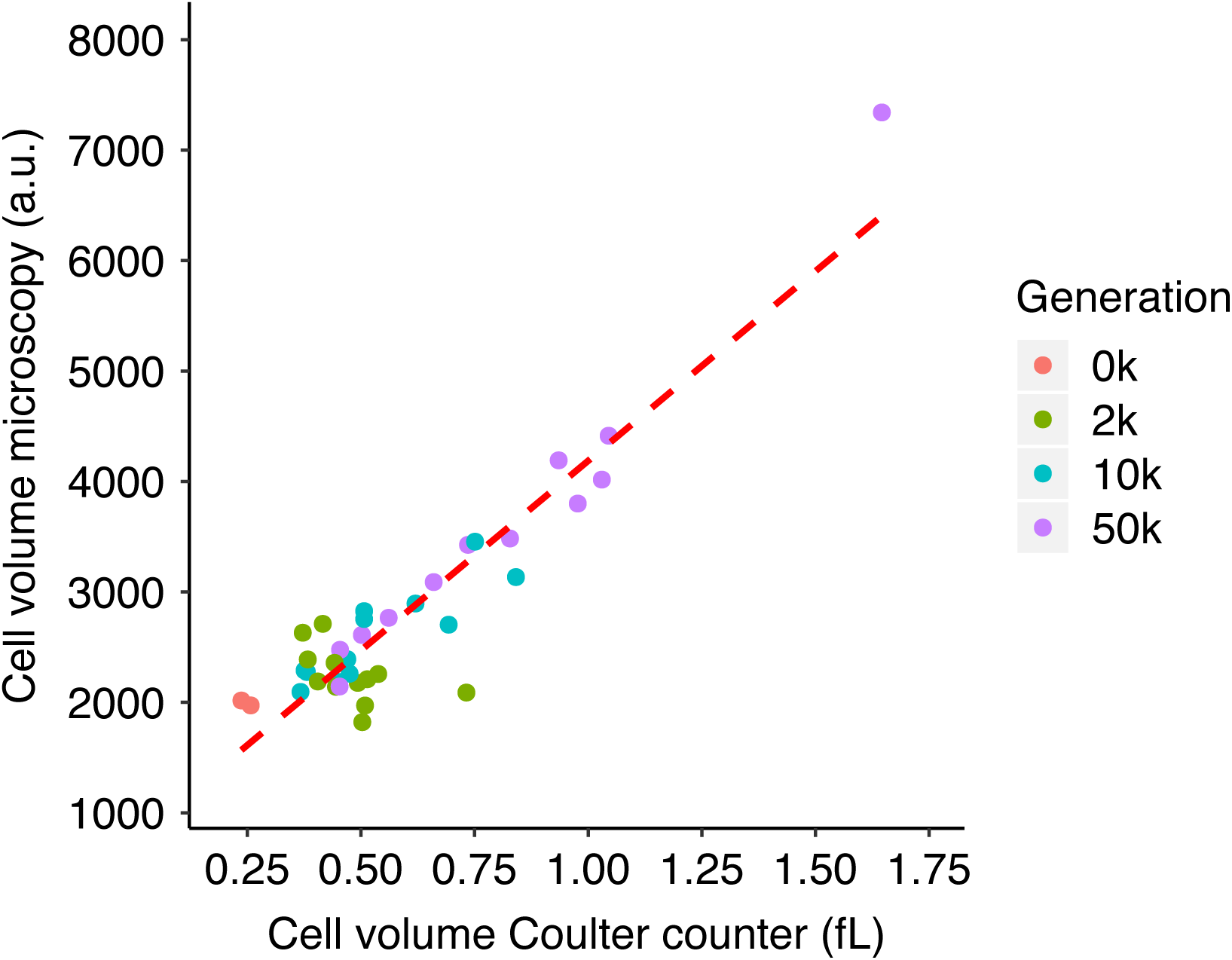
Correlation between cell volume measurements obtained using microscopy and Coulter counter. Volumes obtained by microscopy are expressed in arbitrary units (a.u.) proportional to fL (i.e., µm^3^); volumes obtained using the Coulter counter are expressed in fL. Each point shows the grand median of three assays for clones sampled from the 12 evolving populations or of six assays for the two ancestral strains. Kendall’s coefficient τ = 0.5495, *N* = 38, *p* < 0.0001.

### Temporal trends in cell size in evolved clones

It was previously reported that cell size increased in parallel across all 12 LTEE populations through 10,000 generations, and that the increase in cell volume was strongly correlated with the populations’ improved fitness in the LTEE environment (40). Subsequent papers reported continued fitness gains in the LTEE populations for an additional 40,000 generations (42, 43), albeit at a declining rate of improvement. Here we ask whether cell size also continued to increase, focusing first on the clones isolated from each population at 2,000, 10,000 and 50,000 generations and measured during stationary phase.

Fig. 2A shows that the evolved clones were all larger than their ancestors, although cell size did not always increase monotonically over the course of the LTEE. The median cell volumes of clones sampled from three populations (Ara−2, Ara–3, and Ara–6) were smaller at 10,000 generations than at 2,000 generations. Nonetheless, the median cell volumes of all 12 populations at 50,000 generations were greater than at 10,000 generations. However, the measurement noise associated with the rather small number of biological replicates (i.e., independent cultures) for each clone, and the requirement to correct for multiple hypothesis tests, make it difficult to statistically ascertain the changes in cell volume between clones from successive generations. One possibility is that individual clones are not always representative of the populations from which they were sampled. If that were the case, then we would expect to see more consistent temporal trends in whole-population samples than in clones. We will address that issue in the next section. On balance, the median cell volumes of the evolved clones were on average 1.49, 1.68, and 2.55 times greater than the ancestor at 2,000, 10,000, and 50,000 generations, respectively (one-tailed paired *t*-tests: *p* = 0.0067, 0.0019, and 0.0006, respectively).

**FIG 2.**
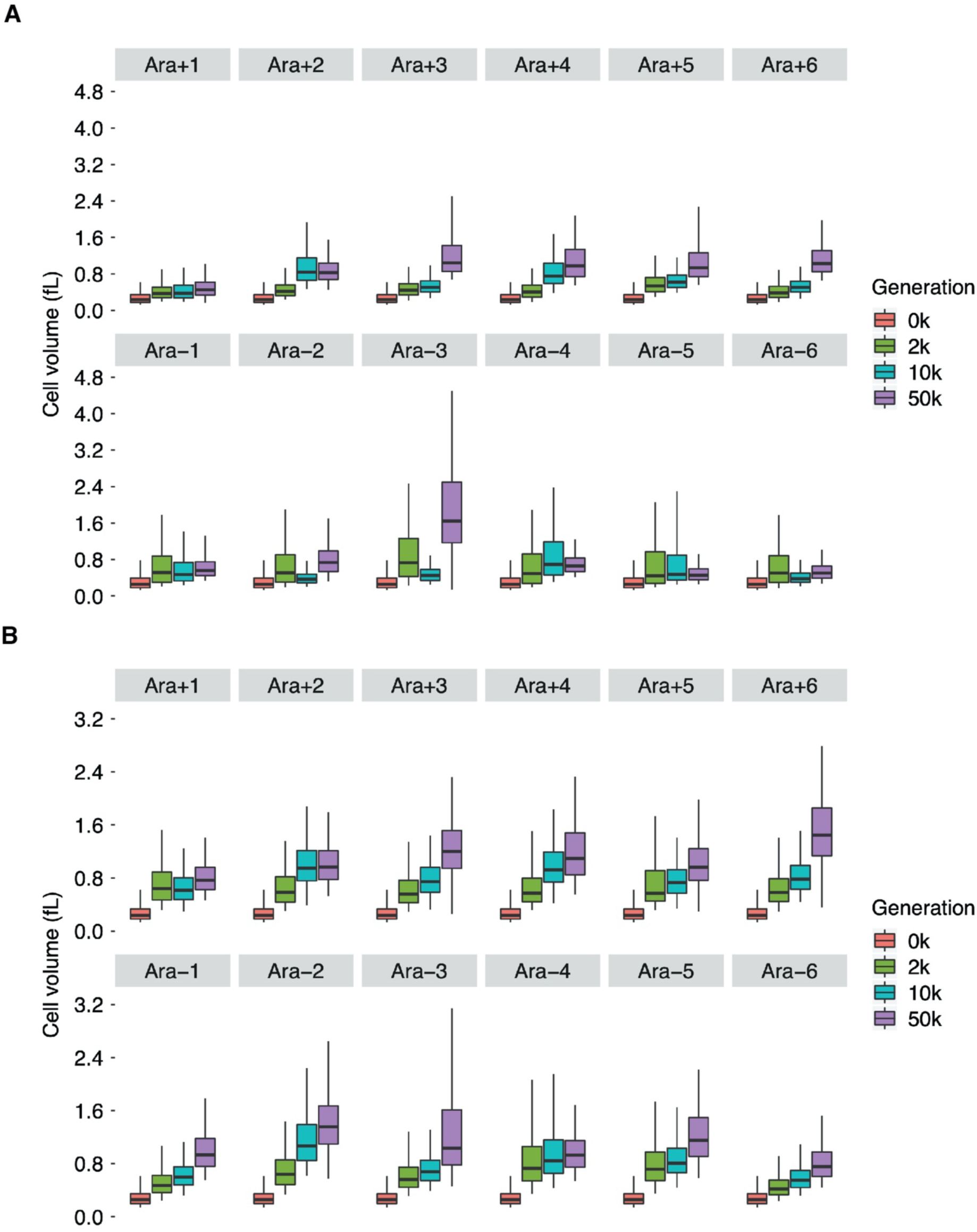
Cell size trajectories for (A) clones and (B) whole-population samples obtained using Coulter counter. Each quantile (5th, 25th, 50th, 75th, and 95th) represents the median of the corresponding quantile from six replicates of each ancestor (REL607 for “Ara+” populations; REL606 for “Ara−” populations) and three replicates for cells sampled from each evolving population.

Besides the possible reversals in median cell size between 2,000 and 10,000 generations in a few populations, two other unusual cases are noteworthy. The 50,000- generation clone from population Ara−3 had by far the largest cells, with a median cell volume that was ∼1.6 times greater than any other population at the same time point (Fig. 2A). That population is the only LTEE population that evolved the capacity to use the abundant citrate in the DM25 medium as an additional carbon source beyond the glucose that limits the other populations (47, 48). The Cit^+^ phenotype is clearly advantageous, although it should also be noted that growth is slower on citrate than on glucose (49). Given that slower-growing *E. coli* cells tend to be smaller than faster-growing cells (14,15, 16, 50), and that this population’s growth shifts in an apparent diauxic manner from glucose to citrate (49), it is surprising that this clone produces the largest stationary-phase cells of any clone we examined. Perhaps these cells are sequestering unused carbon, accounting for their large size; or perhaps the evident stress they face during growth on citrate (49) leads to some decoupling of their growth and division. The other noteworthy population is Ara+1, which showed the smallest increase in cell volume (Fig. 2A). This population also achieved the smallest fitness gains of any of the LTEE populations (42, 43). Given that growth rate is the main determinant of fitness in the LTEE (41), it is therefore interesting (but not surprising) that Ara+1 is both the least fit and produces the smallest cells of any of the LTEE populations.

Fig. 3A compares average cell volumes of the clones between the consecutive generations sampled. These analyses show that average cell size across the 12 LTEE lines increased significantly from the ancestor to generation 2,000, and between 10,000 and 50,000 generations; however, the increase between 2,000 and 10,000 generations was not significant.

**FIG 3.**
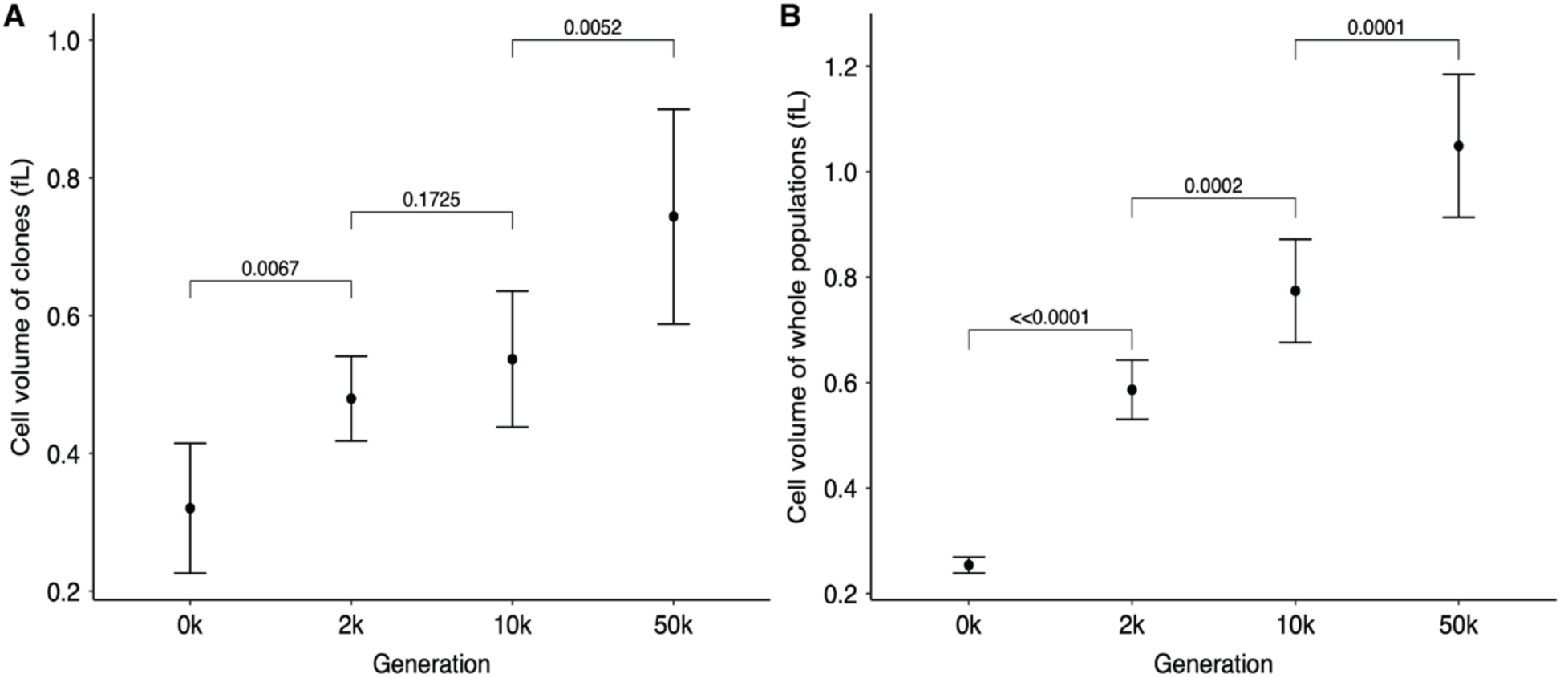
Tests of changes over time in average cell sizes of (A) clones and (B) whole-population samples from the 12 LTEE populations. Each point shows the grand mean of the grand median cell volumes calculated for each population. The 50,000-generation clone from population Ara−3 was an extreme outlier (FIG 2A) and is excluded in panel A; however, the 50,000-generation whole-population sample from this population was not an outlier (FIG 2B). Error bars are 95% confidence intervals, and brackets show the statistical significance (*p* value) based on one-tailed paired *t*-tests. The last comparison in panel A remains significant even if one includes the outlier clone (*p* = 0.0090).

### Monotonic cell size trends among whole populations

We have so far established that the cell volume of clones usually, but not always, increased between the generations tested. However, the evolutionary changes in clones are not always representative of the populations from which they are sampled. For this reason, we measured the cell volumes of whole-population samples at the same three generations to see whether they might show more consistent temporal trends. Fig. 2B shows the cell volume trajectories for these measurements. Indeed, the population samples showed more consistent trends toward larger cells than did the clones. The grand mean trend of the whole populations (Fig. 3B) closely mirrored the overall trend seen for clones (Fig. 3A). However, the correlation between cell volumes measured on clones and whole populations, while highly significant overall, also showed considerable scatter (Fig. S1), indicating that individual clones are not always representative of the populations from where they were sampled. One such difference was that the median volume in the 50,000-generation whole-population sample of Ara–3 was no longer an outlier when compared to the other populations (Fig. 2B), in contrast to the measurements on the individual clones (Fig. 2A). Another difference was the increase in median cell size from the ancestral state to generation 50,000 was much greater in the whole-population sample of Ara+1 than in the individual clone.

Overall, the temporal trend in cell volume does not appear to have reached any upper bound or asymptote, as each generation of whole-population samples that we tested had significantly larger cells than the preceding generation (Fig. 3B). However, the intervals between samples were also progressively longer. Therefore, we calculated the average rate of change in cell volume from the slopes calculated for each population between adjacent time points (Fig. 4). The average rate of cell volume increase was ∼0.17 fL per thousand generations in the first 2,000 generations but dropped to ∼0.02 and ∼0.007 fL per thousand generations in the following 8,000- and 40,000-generation intervals, respectively. In summary, these data show that cell size has continued to increase throughout the long duration of the LTEE, albeit at a decelerating pace and notwithstanding a few atypical evolved clones.

**FIG 4.**
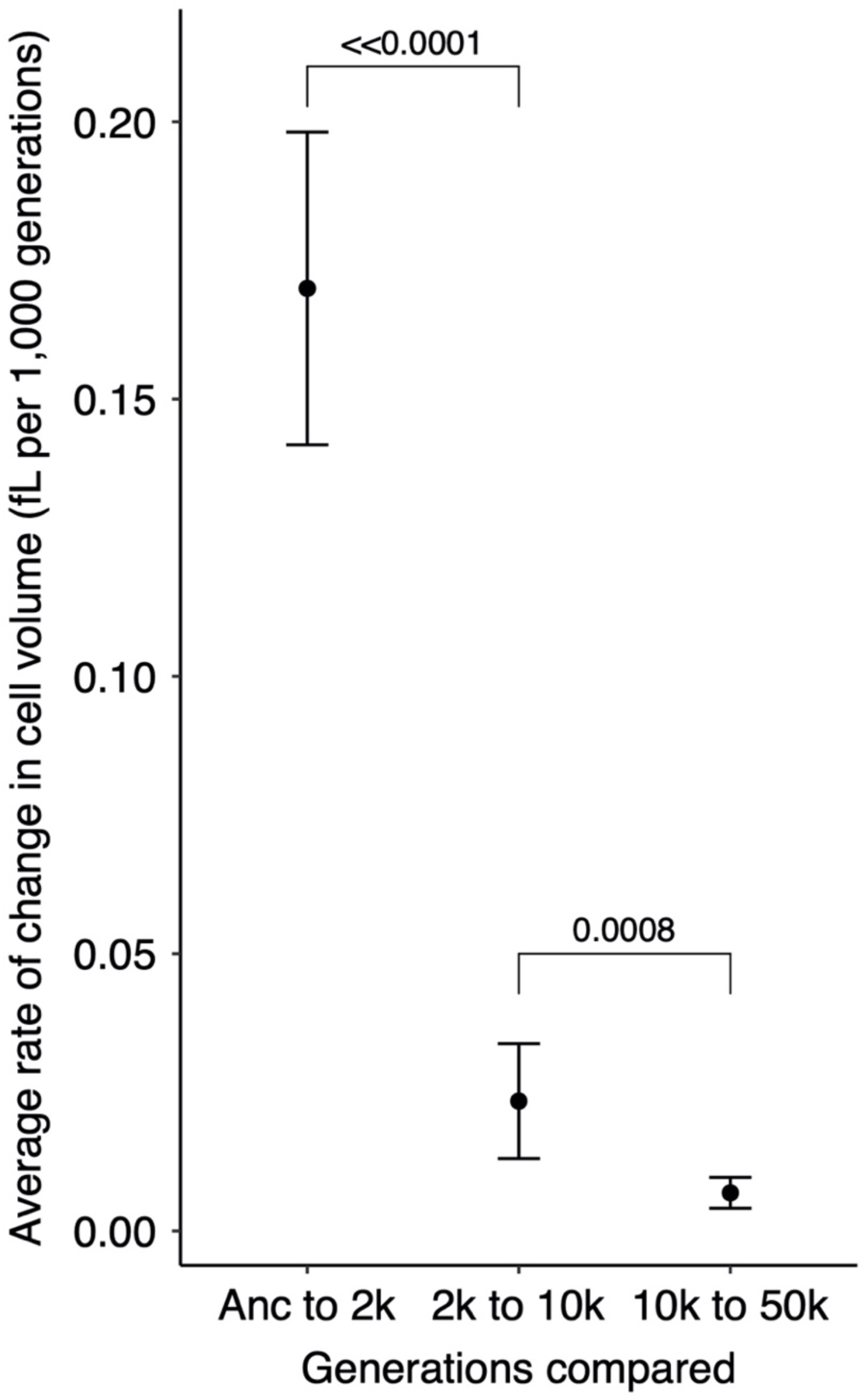
Average rate of cell volume increase. Slopes were calculated for each population over each of three intervals. Each point shows the grand mean for the 12 populations. Error bars are 95% confidence intervals, and brackets show the statistical significance (p value) based on one-tailed Wilcoxon tests, which account for the paired nature of the samples.

### Differences in cell size between exponential and stationary phases

In the sections above, we established the following points: (i) there is good agreement between cell volumes estimated using the Coulter particle counter and by microscopy; (ii) the evolved cells are generally much larger than their ancestors; (iii) there is a nearly monotonic trend over time toward larger cells, although at a declining rate and with a few clones as outliers; and (iv) the independently evolving populations show substantial variation in their average cell sizes after 50,000 generations. All of these conclusions were obtained using cells in stationary phase, and it is of interest to ask whether they also hold for exponentially growing cells. However, examining these issues with exponentially growing cells presents additional challenges. In particular, owing to evolved changes in growth rates and lag times (41), cells from different generations and populations reach mid-exponential-phase growth at different times, complicating efforts to obtain consistent measurements. In addition, the DM25 medium in which the cells evolved is dilute: the stationary-phase population density of the ancestor is only ∼5 x 10^7^ cells per mL, and it is even lower for most evolved clones owing to their larger cells. Hence, cells in mid-exponential-phase growth are usually at densities less than 10^7^ cells per mL. For these reasons, and given the excellent correspondence between Coulter counter and microscopic data, we measured the distribution of cell volumes for exponentially growing cells using only the Coulter counter.

We measured cell volumes of the ancestors and 50,000-generation clones from all 12 LTEE populations 2 h and 24 h after they were transferred into fresh DM25 medium (Fig. 5). At 2h, even the ancestors have begun growing exponentially (41), and none of the evolved strains grow so fast that they would have depleted the limiting glucose by that time (42). The 24-h time point corresponds to when the cells are transferred to fresh medium during the LTEE and hence leave stationary phase. This paired sampling strategy allows us to ask how predictive the stationary-phase cell volumes are of exponentially growing cells. In fact, we found a strong positive correlation in cell volumes measured during exponential growth and stationary phase (Fig. 6). The exponentially growing cells were consistently much larger than those in stationary phase for the ancestors as well as all of the 50,000-generation clones (Fig. 5). For the evolved clones, the volumetric difference as a function of growth phase was ∼2-fold, on average (Fig. S2). It is well known that bacterial cells are larger during exponential growth, with each fast-growing cell typically having multiple copies of the chromosome and many ribosomes to support maximal protein synthesis. In the dilute glucose-limited DM25 minimal medium, cells hit stationary phase abruptly, with the last population doubling using up as much glucose as all the previous doublings combined. The ∼2-fold volumetric difference between the exponentially growing cells and those measured many hours later in stationary phase implies that they typically undergo a reductive division, either as they enter or during stationary phase. At the same time, the range in size between the 12 independently evolved clones was also roughly 2-fold during both growth phases (Fig. 5), which indicates that the striking morphological divergence extends across growth phases.

**FIG 5.**
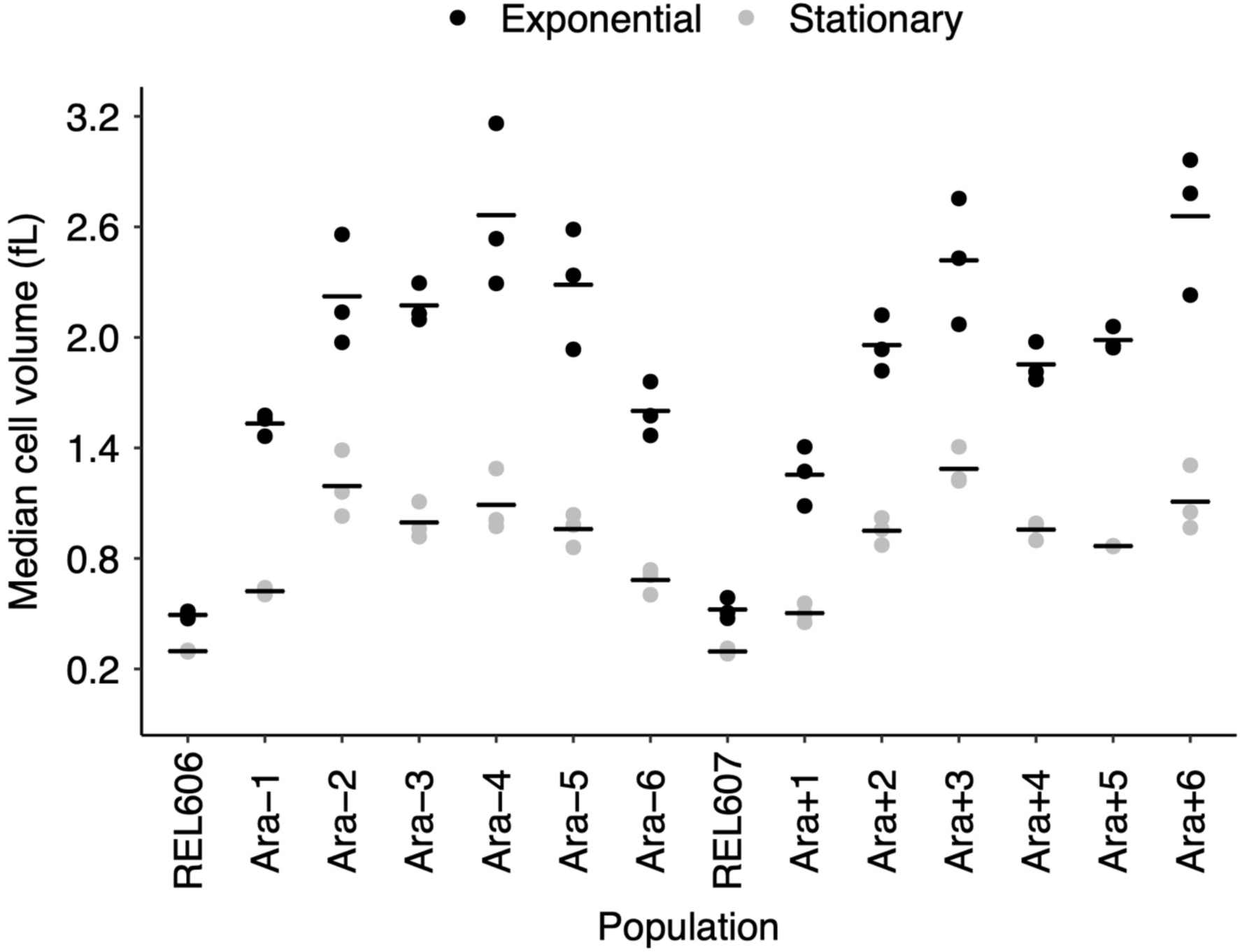
Cell sizes measured during exponential and stationary phases of ancestral strains and 50,000-generation clones from all 12 populations. Each point represents the median cell volume for one assay at either 2 h (exponential growth) or 24 h (stationary phase) in DM25. Horizontal bars are the means of the 3 replicate assays for each strain. The points for some individual replicates are not visible because some values were almost identical.

**FIG 6.**
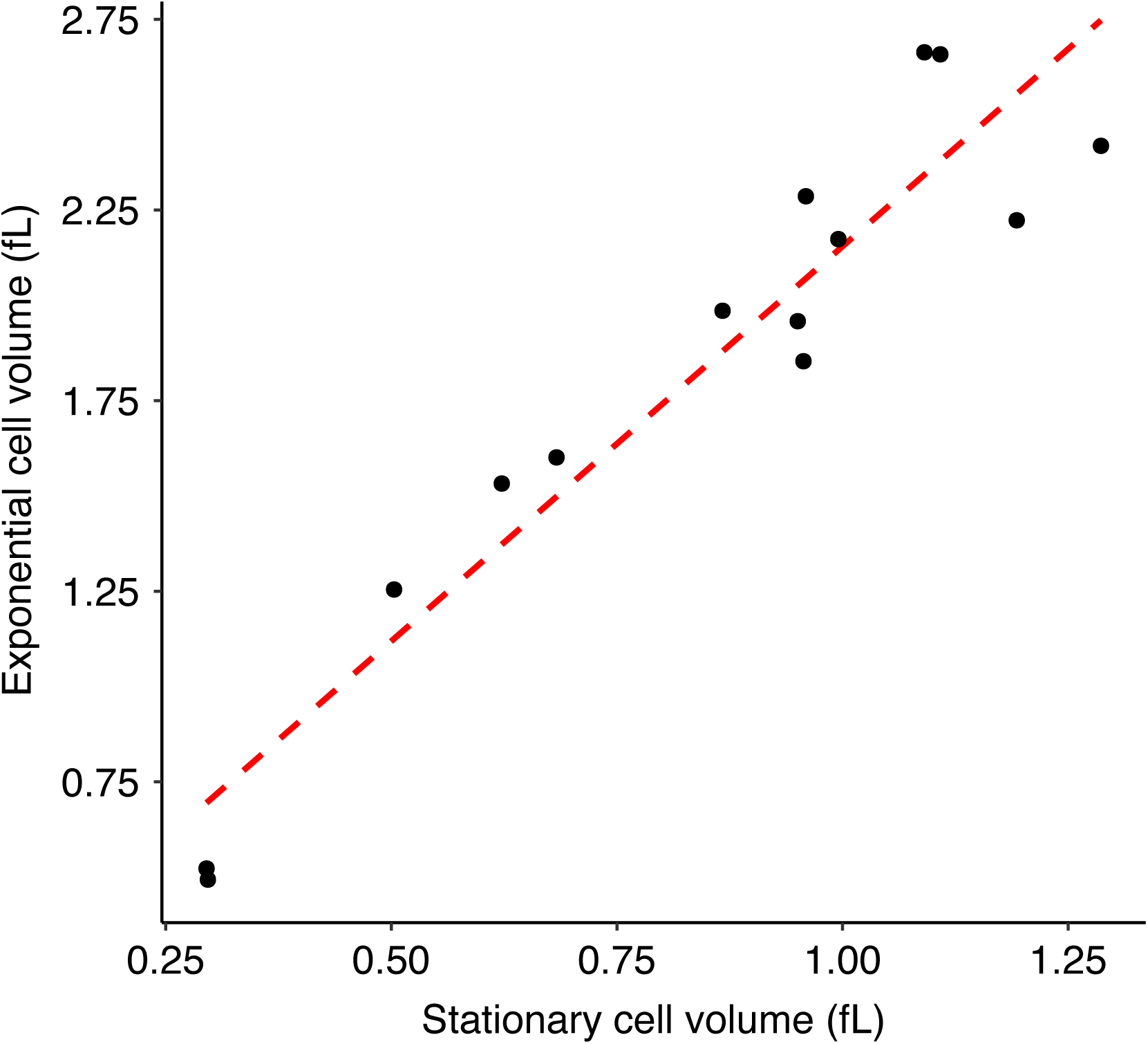
Correlation between cell sizes during exponential growth and in stationary phase. Each point represents the average over 3 replicates of the median cell volume in each growth phase using the data shown in FIG 5. Kendall’s coefficient τ = 0.7582, *N* = 14, *p <*< 0.0001.

### Changes in cell shape

Cell size has clearly increased during the LTEE. Has cell shape also changed? Cell shape has sometimes been regarded as invariant for a given species. For example, *E. coli* has rod-shaped cells that typically maintain an aspect ratio (length-to-width) of ∼4:1, independent of cell volume (51, 31). We examined and analyzed micrographs to see whether the larger cells that evolved in the LTEE maintained their ancestral aspect ratio. Alternatively, larger volumes might have evolved by disproportionate increases in either the length or width of cells. Yet another possibility is that the lineages diverged in their aspect ratios not only from their common ancestor, but also from one another. Fig. 7 shows representative micrographs of the ancestors and the 50,000-generation clones. It is readily apparent that the different lineages have evolved different aspect ratios. To investigate these differences more systematically, we processed multiple micrographs of the ancestors and clones from generations 2,000, 10,000, and 50,000 using the *SuperSegger* package (52). Across all of the samples in total, we obtained lengths and widths (cross-sectional diameters) from >87,000 cells (see Methods). As a reminder, an increase in the aspect ratio relative to the ancestor implies a higher SA/V ratio for a given volume, whereas a decline in the aspect ratio indicates the opposite. Of course, having a larger cell alone also reduces the SA/V ratio, even without a change in the aspect ratio. One would typically expect a greater SA/V ratio to be beneficial for resource acquisition, and therefore we might expect the evolved clones to have higher aspect ratios than the ancestral strains, especially given their increased volumes.

**FIG 7.**
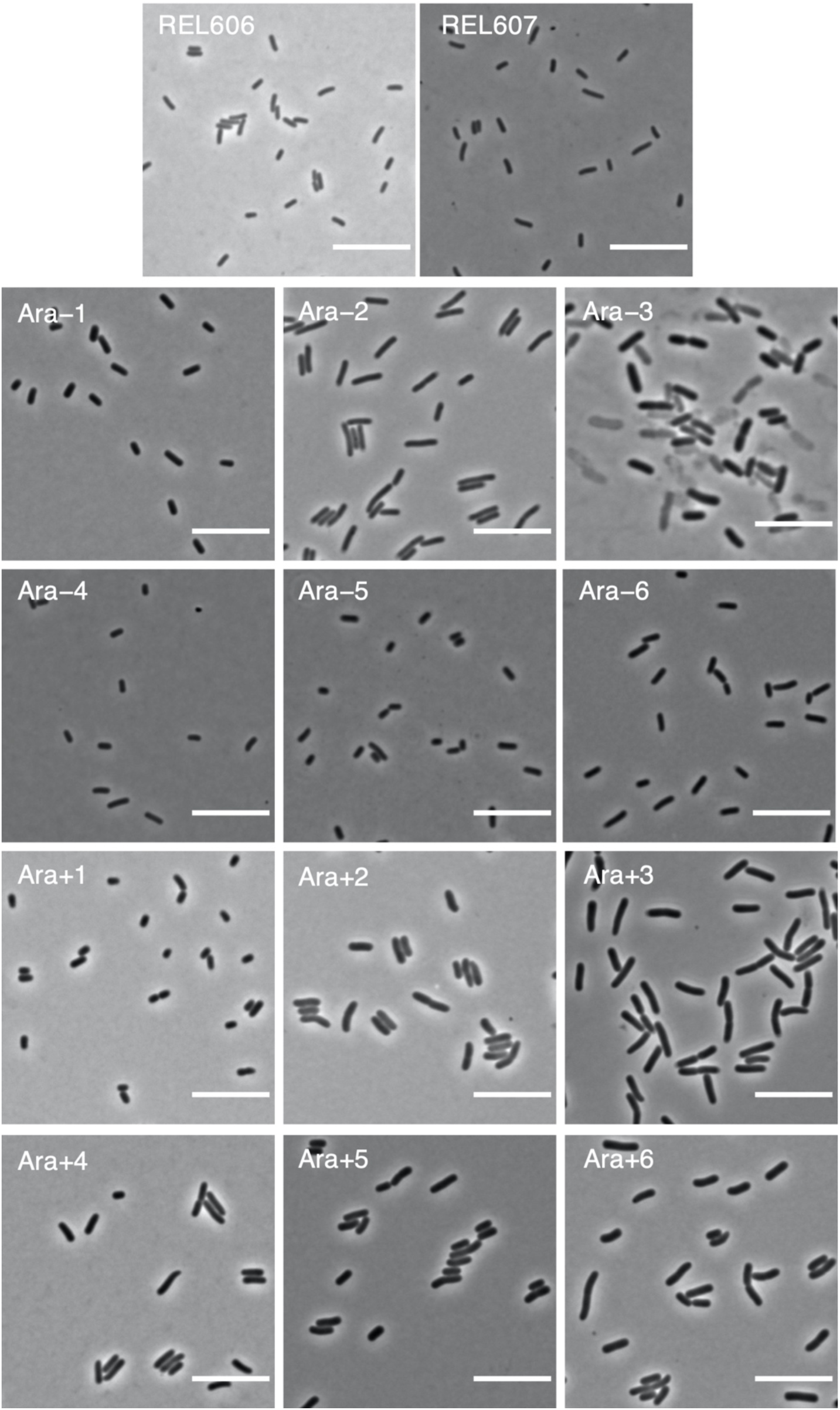
Representative micrographs of ancestors (REL606 and REL607) and evolved clones from each population at 50,000 generations. Phase-contrast images were taken at 100 x magnification. Scale bars are 10 µm.

In fact, however, the opposite trend held, at least for the first 10,000 generations, as shown in Fig. 8. Clones from 10 of the 12 populations, at both 2,000 and 10,000 generations, tended to produce relatively wider than longer cells in comparison to the ancestor (*p* = 0.0386 based on a two-tailed sign test at each time point). By 50,000 generations, the clones were split evenly: 5 had aspect ratios greater than the ancestor, 5 had aspect ratios lower than the ancestor, and 2 had aspect ratios nearly identical to the ancestor. Note that the 50,000-generation clone from population Ara–3 is an extreme outlier, with cells that are exceptionally long and very large. This population is the one that evolved the novel ability to grow on citrate (47, 48, 49), and its unusual morphology is presumably related to its distinct metabolism.

**FIG 8.**
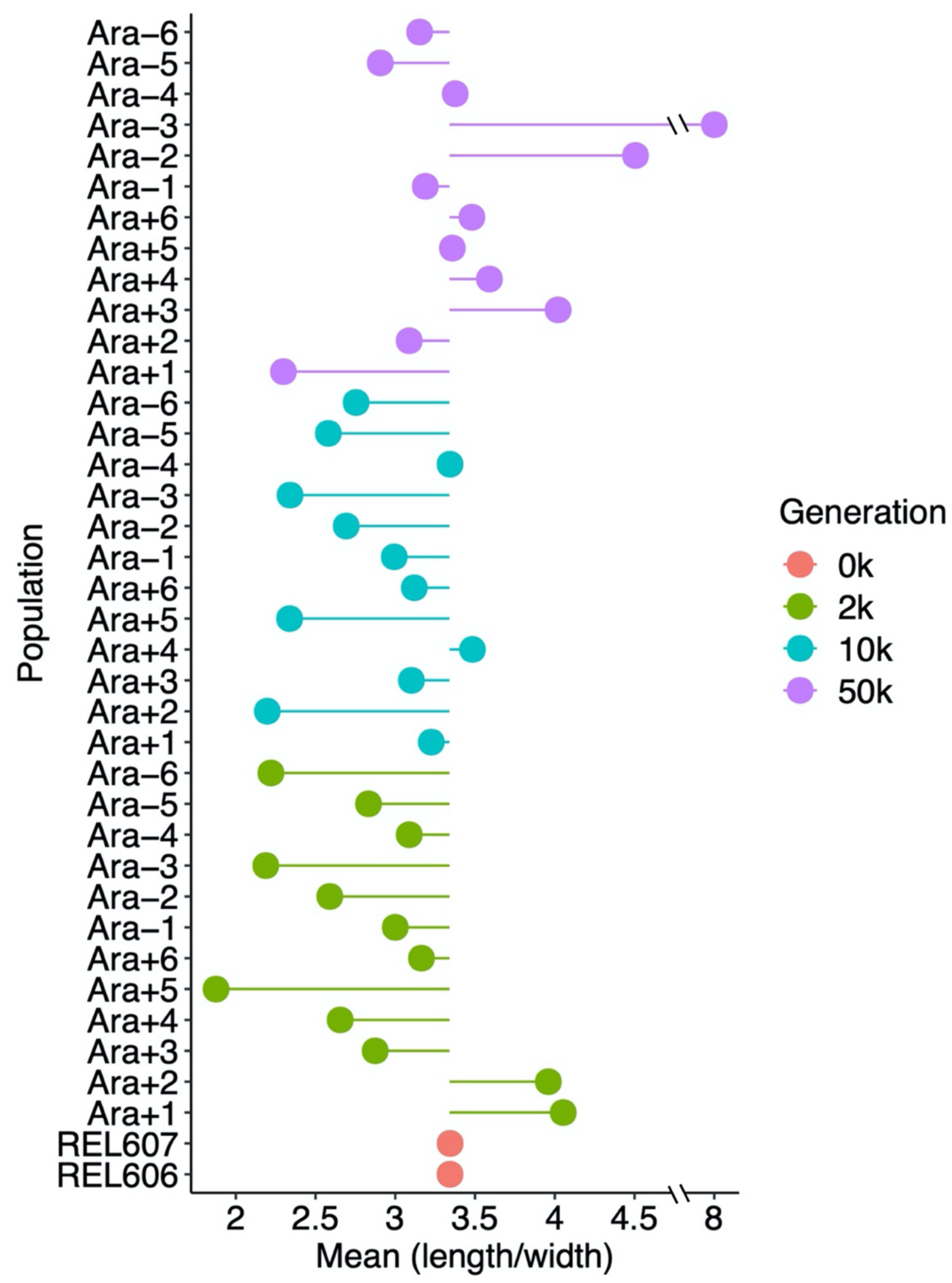
Average cell aspect ratios (length/width) of ancestral and evolved clones. Each point shows the mean ratio for the indicated sample. The lines show deviations in the aspect ratio from the ancestral state. The mean aspect ratios were calculated from three replicate assays in all but 4 cases (Ara−4 at 10,000 generations; Ara−2, Ara−4, and Ara−5 at 50,000 generations), which had two replicates each.

Fig. 9A shows the average length-to-width ratios and their associated 95% confidence intervals, excluding the Cit^+^ outlier at 50,000 generations. The ancestral cells had an average length-to-width ratio of 3.37. Recall that *E. coli* has been reported to typically maintain an average aspect ratio of about 4:1 (33, 51, 53). The aspect ratio we see is somewhat smaller. This difference might reflect variation between strains (the LTEE ancestor is a derivative of *E. coli* B, not K12) or some other factor. In any case, the mean aspect ratio across the evolved lines declined to 2.90 and 2.87 at 2,000 and 10,000 generations, respectively, and then increased to 3.39 at generation 50,000, almost identical to the ancestral ratio. The early decline in the aspect ratio is statistically significant, as is the subsequent reversal (Fig. 9A). This reversal would increase the SA/V ratio somewhat. However, it might not be sufficient to offset the reduction in the SA/V ratio associated with the much larger cell volumes at 50,000 generations. On balance, the LTEE lines evolved larger cell volumes by first increasing disproportionately in width, and later increasing their length, possibly to the benefit of a somewhat more favorable SA/V ratio.

**FIG. 9.**
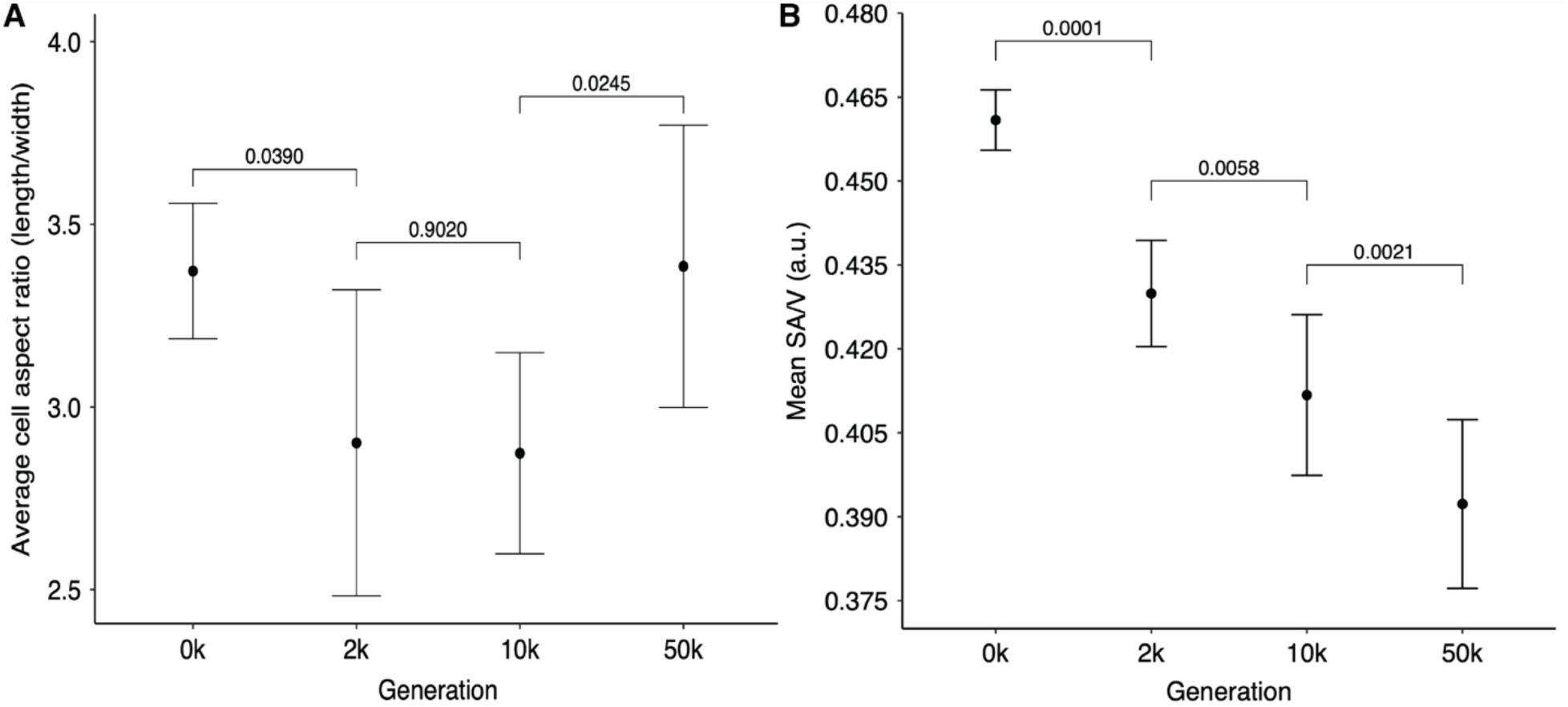
Tests of changes over time in cell aspect and surface-to-volume (SA/V) ratios. (A) Evolutionary reversal of cell aspect ratio. Each point is the grand mean of the cell aspect ratio (length/width) for the ancestors and evolved clones. N = 12, except at 50,000 generations, where N = 11 after excluding the outlier clone from the Ara−3 population. Errors bars are 95% confidence intervals, and brackets show the statistical significance (p value) based on two-tailed *t*-tests. The tests were paired for clones sampled from the same population at the consecutive time points, and the Ara−3 population was excluded from the final test. (B) Monotonic decline in SA/V ratio over 50,000 generations. Each point shows the grand mean of the average ratio calculated for the ancestor and evolved clones. Error bars are 95% confidence intervals, and brackets show the statistical significance (*p* value) based on one-tailed paired *t*-tests.

### Analysis of changes in the SA/V ratio

The reversion of the evolved clones to their ancestral aspect ratio (Fig. 9A), coupled with their overall increase in cell volume (Fig. 3A), raises the question of how much their SA/V ratios have changed. If selection to increase the diffusion of nutrients into cells is strong in the LTEE, then increasing cell length would be beneficial. However, the larger cell volume would have the opposite effect. To examine the net result of these changes, we calculated the SA/V ratio of the evolved clones using the equations for spherocylindrical cells from Ojkic et al. (32), which we presented in Introduction. We used the length and width values measured for clones using *SuperSegger* to compute for each cell *γ*, which depends on the aspect ratio, and from that the cell’s surface area. We then divided that value by the cell’s estimated volume to obtain its SA/V ratio. Given the early trend toward wider cells (lower aspect ratios) and the larger cell volumes at later generations, we expected lower SA/V ratios for the evolved clones relative to the ancestors, despite the later reversion toward the ancestral aspect ratio. Indeed, all 36 evolved clones had a SA/V ratio that was lower than the ancestors (Fig. 10).

**FIG 10.**
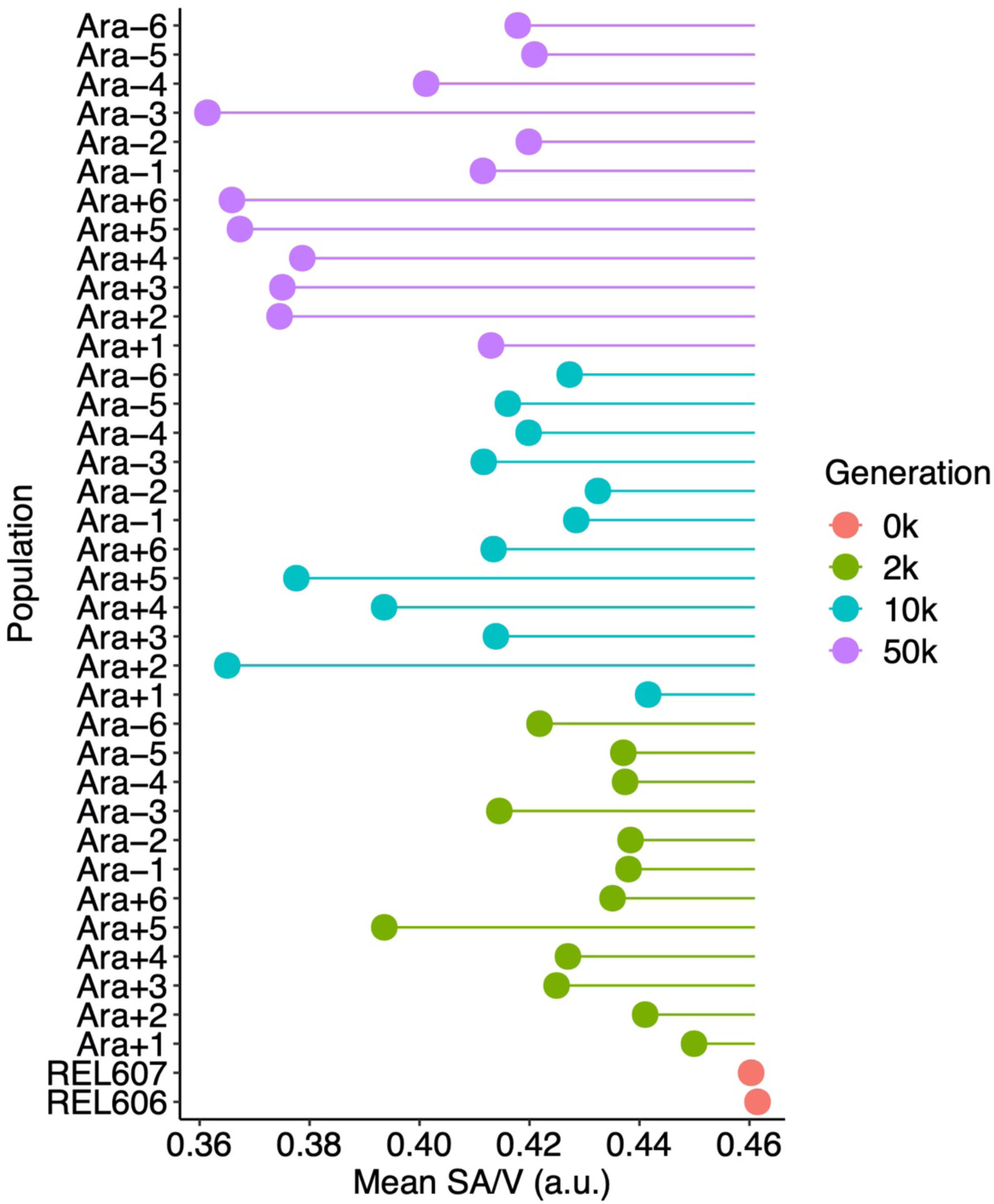
Average surface area-to-volume ratio (SA/V) of ancestral and evolved clones. The surface area and volume of individual cells were calculated from microscopic images, as described in the text, and their ratio has arbitrary units (a.u.) proportional to µm^−1^. Each point shows the mean ratio for the indicated sample. The lines show deviations in the ratio from the ancestral state. The means were calculated from three replicate assays in all but 4 cases (Ara−4 at 10,000 generations; Ara−2, Ara−4, and Ara−5 at 50,000 generations), which had two replicates each.

Fig. 9B shows the average SA/V ratio and associated 95% confidence intervals over time. We included the 50,000-generation Ara−3 clone in this analysis because its SA/V ratio (Fig. 10), unlike its aspect ratio (Fig. 8), was not an extreme outlier; that is, its atypical aspect ratio was largely offset by its large average cell volume (Fig. 2A). The mean SA/V ratio declined monotonically and significantly from 0.461 in the ancestor to 0.430, 0.412, and 0.392 at 2,000, 10,000, and 50,000 generations, respectively. Even the reversion to the ancestral cell aspect ratio between 10,000 and 50,000 generations (Fig. 9A) was insufficient to offset the increase in cell volume over that same interval (Fig. 3A).

We also performed an isometric analysis to assess the extent to which the reversion to the ancestral aspect ratio between 10,000 and 50,000 generations changed the SA/V ratio. To do so, we used the cell aspect ratios measured at 10,000 generations and compared the average SA/V ratio at 50,000 generations to the hypothetical average using the earlier aspect ratios. The average SA/V ratio at 50,000 generations was ∼6% higher as a consequence of the change in cell aspect ratio (Fig. S3), and this difference was significant (*p* = 0.0144). Even so, the mean SA/V ratio continued to decline (Fig. 9B) because the change in average cell aspect ratio over this period (Fig. 9A) was insufficient to offset the increase in average cell volume (Fig. 3A).

### Nearly spherical cells in one LTEE population

While examining micrographs, we observed that cells from the Ara+5 population at 2,000 and 10,000 generations looked like stubby rods, many of which were almost spherical (Fig. 11). By 50,000 generations, however, the cells were rod-shaped (Fig. 7), suggesting that one or more mutations in morphogenic genes might contribute to this phenotype.

**FIG 11.**
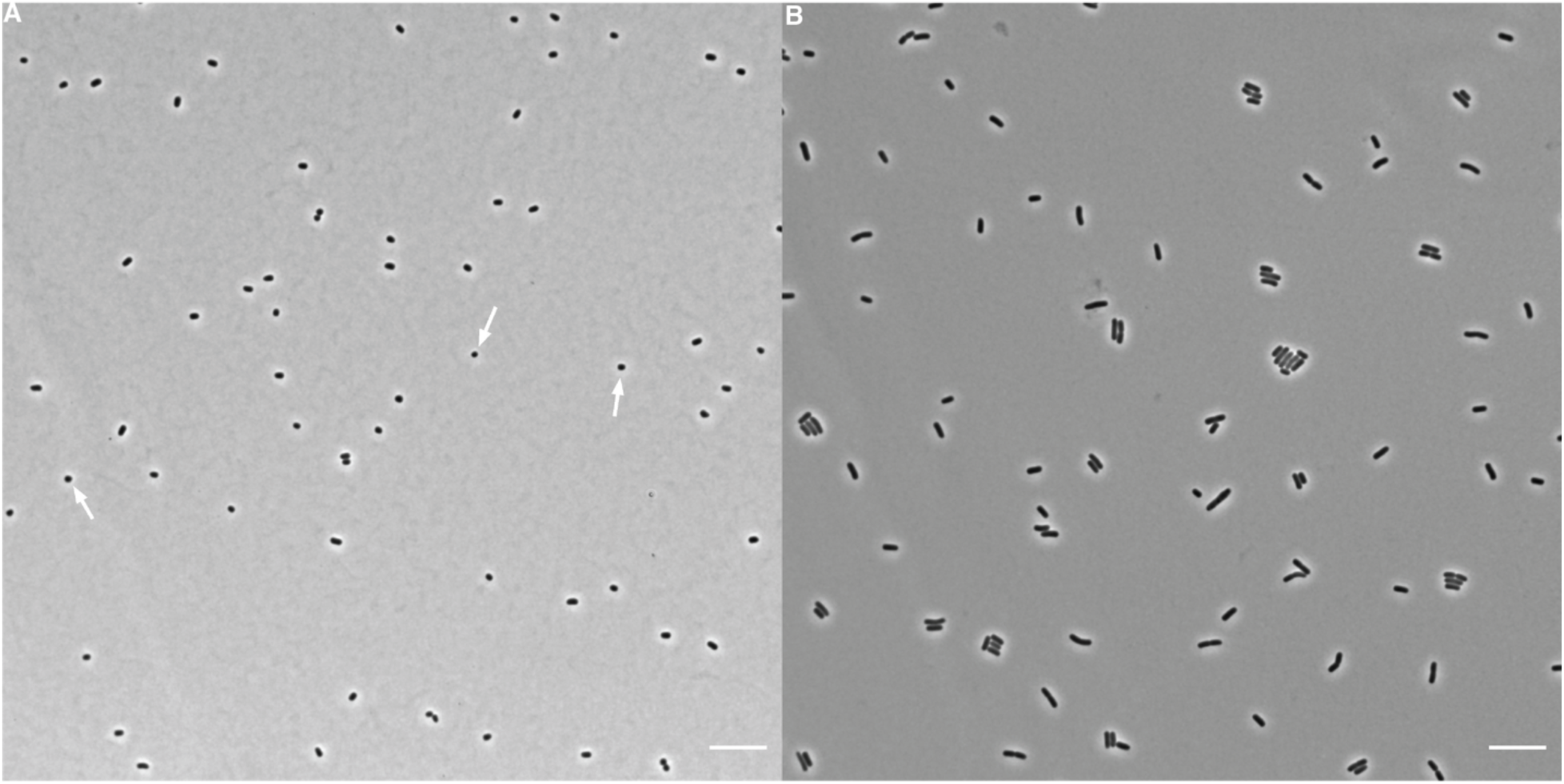
Representative micrographs of cells from (A) 2,000-generation and (B) 50,000- generation clones of the Ara+5 population. Phase contrast images were taken on an inverted microscope at 100 x magnification. Scale bars are 10 µm. Arrows point to examples of nearly spherical cells in the earlier sample, which are not seen in the later one.

The typical rod-shaped cell morphology in *E. coli* is maintained by several proteins including MreB, MreC, MreD, MrdA (PBP2), and MrdB (RodA) (46, 54). To this end, we examined published whole-genome sequence data (55) for the clones in our study to identify any mutations in these genes. By 50,000 generations, all but one of the 12 lines (Ara−5) had nonsynonymous mutations in at least one of these five shape-maintaining genes (Fig. 12). There were also a few synonymous changes, which were seen only in populations that had evolved point-mutation hypermutability, as well as one indel. However, the majority of mutations that arose and reached high frequency in these genes were nonsynonymous changes.

**FIG 12.**
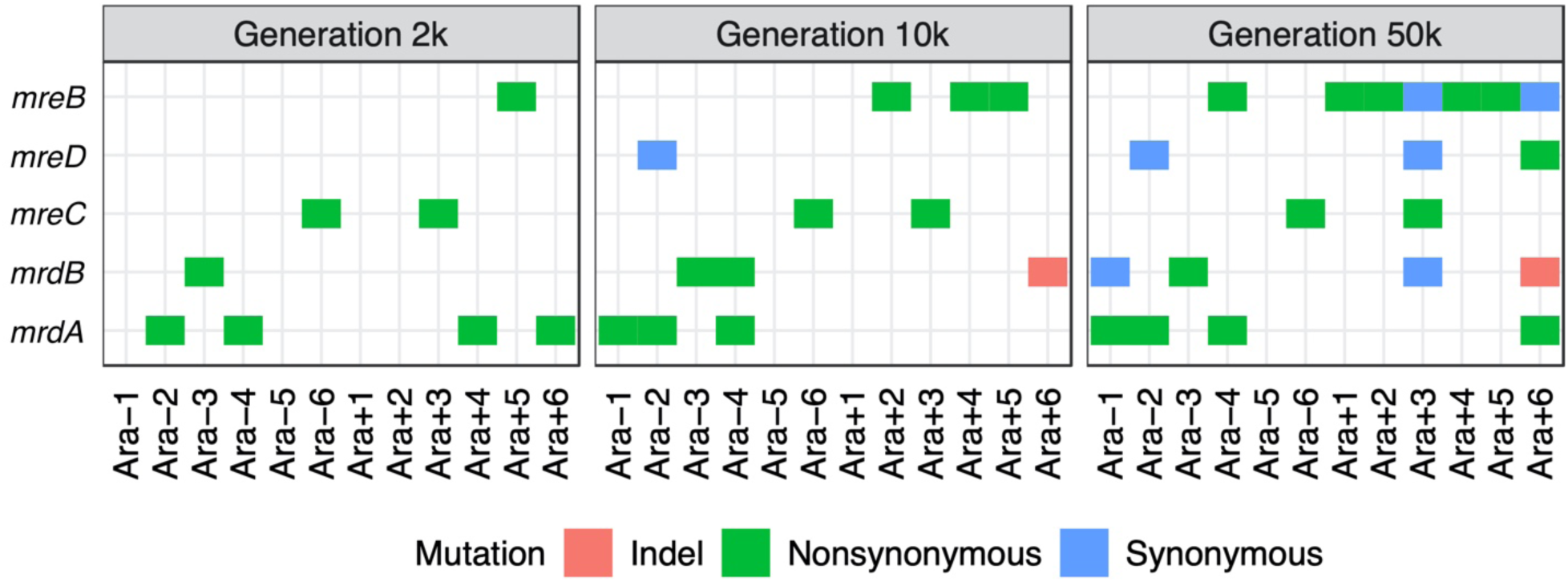
Parallel mutations in genes known to be involved in the maintenance of rod-shaped genes. Nonsynonymous mutations were found in all populations except Ara−5 by 50,000 generations. Populations Ara−2, Ara−4, Ara+3, and Ara+6 evolved hypermutable phenotypes between generations 2,000 and 10,000; populations Ara−1 and Ara−3 did so between 10,000 and 50,000 generations. Hence, all synonymous mutations were found in lineages with a history of elevated point-mutation rates.

The 2000-generation clone from the Ara+5 population that produced the stubby cells had a single nonsynonymous mutation in *mreB*. This mutation was also present in the clones sampled from this population at 10,000 and 50,000 generations. There were no other mutations in the other four rod-shape maintenance genes at any of the timepoints. *E.* coli cells have been shown to become spherical when MreB is depleted (54), which strongly suggests that the *mreB* mutation is responsible for the stubby morphology observed in the early generations of this population. The fact that the Ara+5 cells were not stubby at 50,000 generations, despite the *mreB* mutation, suggests some compensatory change that did not involve the five morphogenic genes considered here. Four other populations also had nonsynonymous *mreB* mutations by generation 50,000 (Fig. 12). Of these four, the clone from population Ara+1 also produced rather stubby cells (Fig. 7), resulting in the lowest aspect ratio of any of the 50,000-generation clones (Fig. 8). Whether the diverse effects of the *mreB* mutations on cell shape reflect the different mutations, the genetic backgrounds on which they arose, or both remains to be determined.

### Cell volume and fitness have remained highly correlated in the LTEE

Cell size and relative fitness were previously shown to be strongly correlated during the first 10,000 generations of the LTEE (40). The fitness of these populations has continued to increase throughout this experiment (42, 43). In light of the continued increase in cell volumes reported in this work, we expected that cell size and fitness would continue to be correlated. To test this, we used the relative fitness data previously collected for the 12 LTEE populations through 50,000 generations (42), and we asked how well those fitness values correlate with the cell volumes we measured for the ancestors and the whole-population samples from three later generations. Fig. 13 shows that cell volume and relative fitness have remained significantly correlated, although with substantial scatter. Some of this scatter reflects increased measurement noise when estimating relative fitness in later generations. These estimates are obtained by competing the evolved populations against a marked ancestor; as the relative fitness of the evolved bacteria increases, it becomes more difficult to enumerate accurately the relative performance of the two competitors.

**FIG 13.**
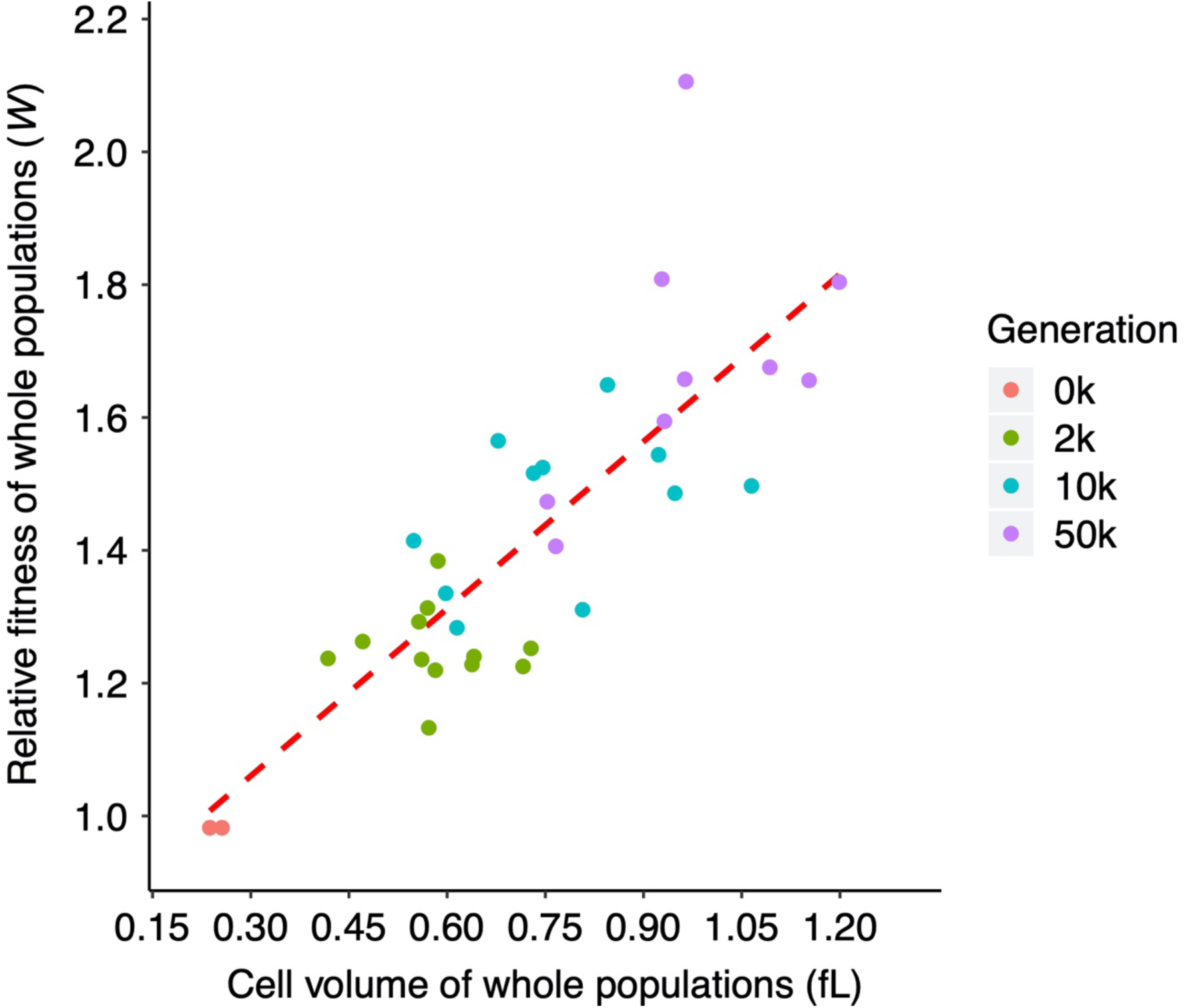
Correlation between mean fitness relative to the LTEE ancestor and grand median cell volumes, both based on whole-population samples. Four points (Ara+6 at 10,000 generations; Ara−2, Ara−3, and Ara+6 at 50,000 generations) are absent due to missing fitness values reported by Wiser et al. (2013). Kendall’s τ = 0.6066, *N* = 34, *p* < 0.0001.

### Elevated cell mortality in the population that evolved to grow on citrate

We observed what we call “ghost” cells in micrographs of the 50,000-generation Cit^+^ clone from the Ara−3 population. These cells were quite distinct from the ancestral strain and evolved clones from all other populations (Fig. 7). In terms of contrast with their background, the ancestor and other evolved clones had uniformly dark and opaque cells, in contrast to the lighter agar pad on which they were placed for imaging. Many of the Cit^+^ cells, by comparison, were translucent (Fig. 7). Most translucent cells appeared intact, although we also saw some fragmented cells. We presume that the translucent cells that appear intact are nonetheless either dead or dying.

We also grew the Cit^+^ clone in DM0, which is the same medium as used in the LTEE and our other experiments, except DM0 contains only the citrate but no glucose. The proportion of ghost cells is even higher in this citrate-only medium (Fig. 14). Some translucent cells had small punctations, or dots, within the cytoplasm, often at the cell poles (Fig. 14). These dots are reminiscent of the polyhydroxyalkanoate storage granules that some bacterial species produce under conditions where their growth is unbalanced (56, 57) or when cells are otherwise stressed (58, 59). It is also possible that these dots comprise the nucleoid or some other remnant of a leaky cytoplasm.

**FIG 14.**
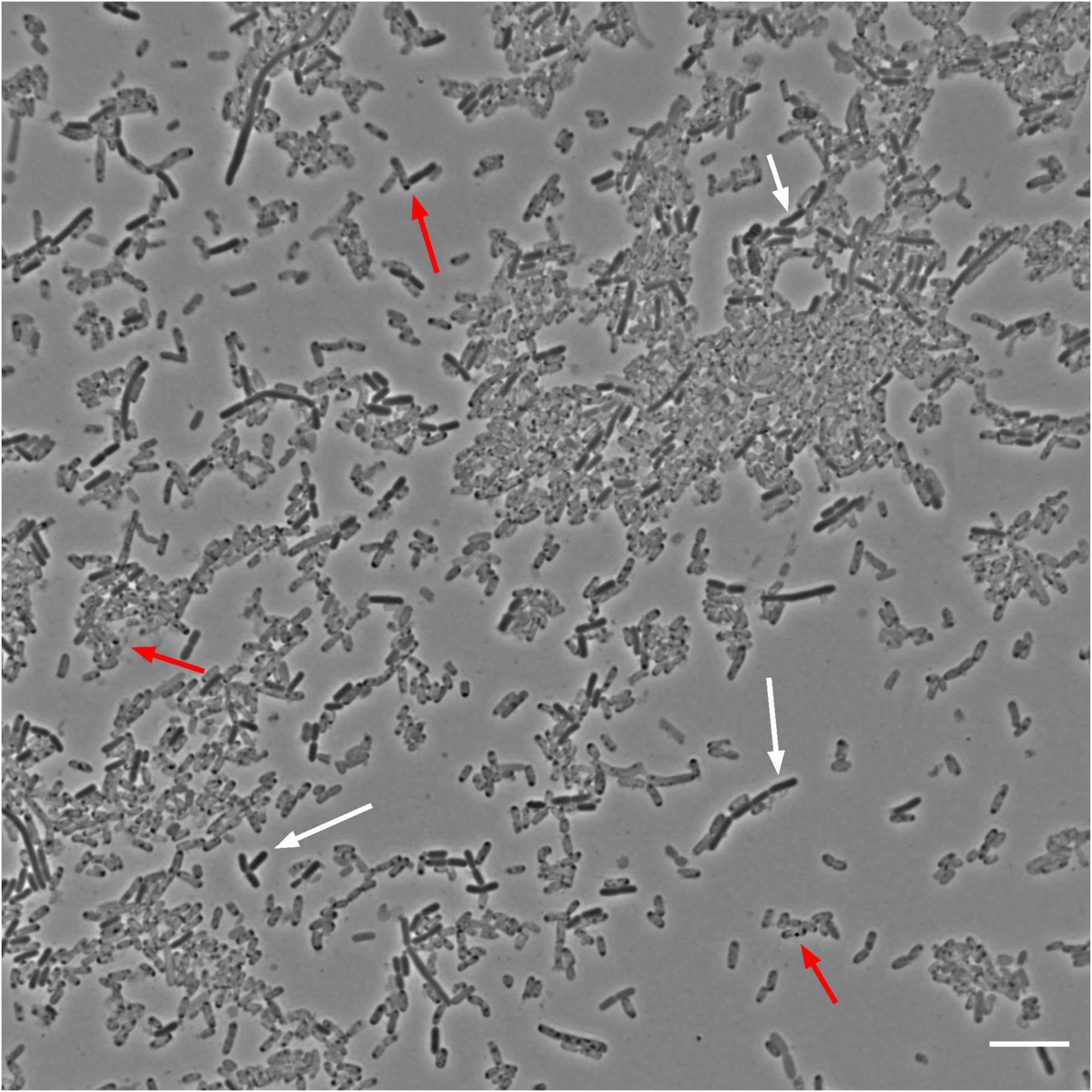
Representative micrograph of 50,000-generation Cit^+^ clone from population Ara−3 grown in DM0. As shown in FIG 7, we observed translucent “ghost” cells in the only population that evolved the capacity to use citrate in the LTEE medium (DM25). This clone can also grow on citrate alone in the same medium except without glucose (DM0), which increased the proportion of presumably dead or dying ghost cells. Red arrows point to several ghost cells, some of which have darker punctate inclusions; white arrows point to several more typically opaque and presumably viable cells. Scale bar is 10 µm.

It is noteworthy that we observed these anomalous ghost cells at any appreciable frequency only in the unique Cit^+^ population (47, 48). This observation of ghost cells, and the implication that many cells in this population are dead or dying, is supported by other observations that indicate the Cit^+^ cells struggle with maintaining balanced growth on citrate (49). To test whether the ghost cells are dead, dying, or at least physiologically incapacitated, we labeled stationary-phase cultures using a two-color live/dead stain. Our methods, full results, and in-depth analyses of these labelling experiments are presented elsewhere (49). Here we present a subset of the data, with an analysis that specifically compares the ancestor (REL606) and 50,000-generation Cit^+^ clone (REL11364). Fig. 15A shows representative micrographs of the ancestral and evolved Cit^+^ cells grown to stationary phase in the standard DM25 medium that contains glucose as well as citrate. Fig. 15B shows the estimated proportions of live (green) and dead (red) cells, obtained by pooling data from 5 independent cultures (i.e., biological replicates) for each clone. There was much more cell death in the cultures of the Cit^+^ clone when compared to the ancestor. On average, 43.6% of the Cit^+^ cells were scored as dead, based on greater intensity of the corresponding dye. By contrast, only 13.2% of the ancestral cells were scored as dead, and they exhibited much weaker intensity of that dye (Fig. 15A). The difference in the proportion of dead cells between the ancestor and the Cit^+^ clone is highly significant (*t* = 2.9304, df = 8, one-tailed *p* = 0.0094). This result thus supports our hypothesis that the ghost cells seen in our original micrographs of the Cit^+^ clones were indeed dead or dying.

**FIG 15.**
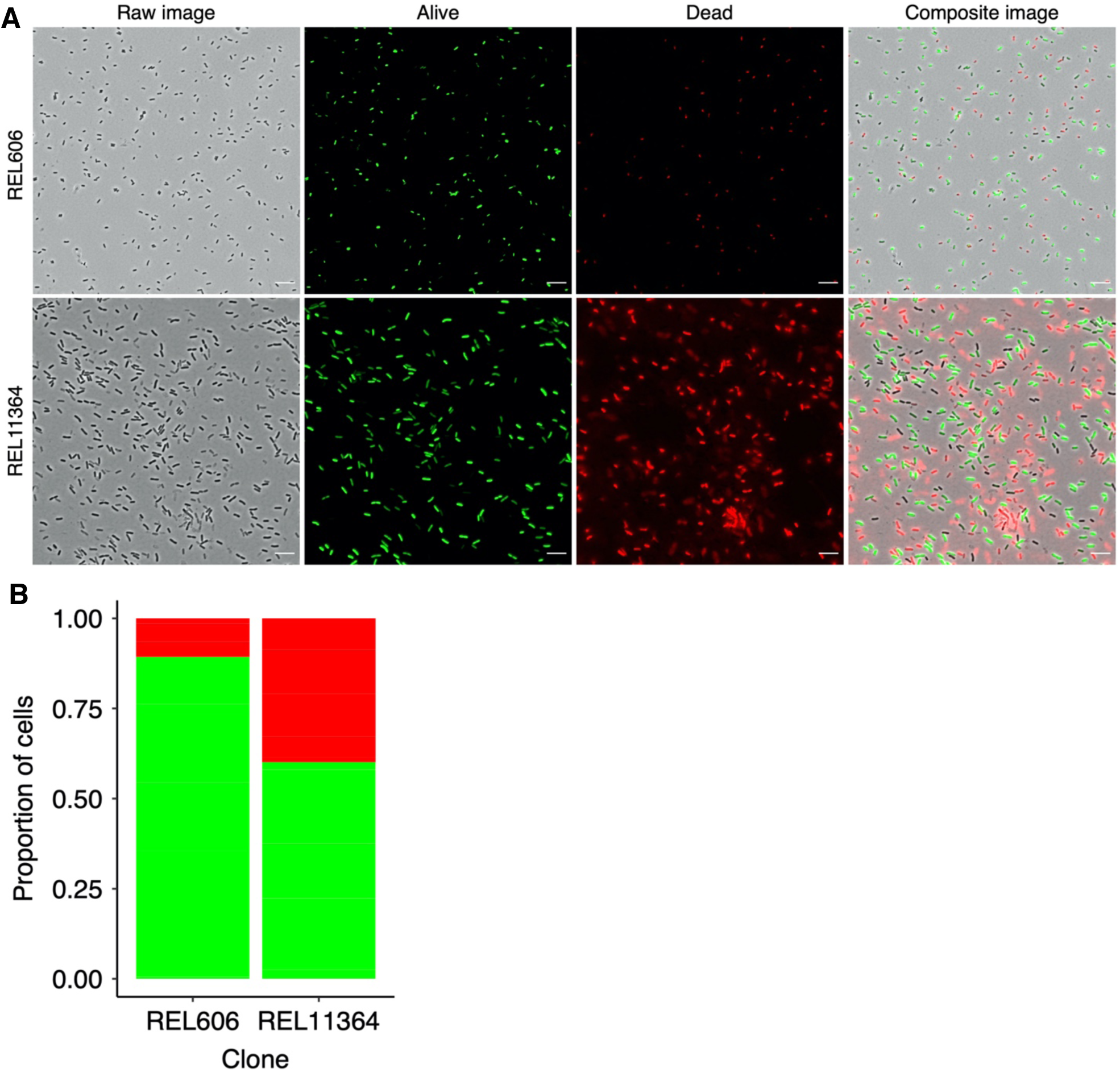
Comparison of cell death in the ancestor and Cit^+^ clone. (A) Representative micrographs showing live-dead staining of the LTEE ancestor (REL606) and the 50,000- generation Cit^+^ clone from population Ara−3 (REL11364), both grown in DM25. Scale bars are 10 µm. (B) Proportions of cells scored as alive (green) or dead (red), based on two-color stain assay. For each clone, we assayed cells from 5 biological replicates, which have been pooled in this figure.

## Discussion

During the first 10,000 generations of the LTEE, 12 populations of *E. coli* increased in fitness and cell size as they evolved in and adapted to their glucose-limited minimal medium (40). The increase in cell size was unexpected, given the fact that larger cells have greater metabolic demands and have SA/V ratios that are less favorable for supporting those demands. In the >60,000 generations since that study, the populations have continued to adapt to the glucose media, and their fitness has continued to increase with trajectories that are well described by a simple power law (42, 43). In this study, we sought to determine if cell size has continued to increase, and whether cell size still correlates with fitness. We measured changes in cell volume and shape for clones and whole-population samples. We used two methods: a Coulter counter that directly measures cell volume, and microscopy that allowed us to analyze both cell volume and shape using machine learning. The average cell volumes measured using the two methods were well correlated (Fig. 1).

The average cell increased monotonically over time in the whole-population samples (Figs. 4−5). Clones from three populations (Ara−2, Ara−3, Ara−6) deviated from this monotonic trend, producing smaller cells at 10,000 than at 2,000 generations (Fig. 2A). These idiosyncratic cases implies within-population heterogeneity. They might also be due, in part, to the clones being studied in an environmental context different from that in which they evolved. As an indication of the relevance of both of these explanations, two ecologically and genetically distinct lineages have coexisted in the Ara−2 population since ∼6,000 generations, with coexistence mediated by differential growth on glucose and acetate, a metabolic byproduct (60, 61). In any case, our data show that average cell size and mean fitness have remained significantly correlated in the LTEE through 50,000 generations (Fig. 13), despite variation within and between populations.

We obtained most of our data on average cell size with cells in stationary phase, at the end of the LTEE’s standard 24-hour period prior to the transfer into fresh medium. We did so because analyzing exponentially growing cells presents additional challenges. In particular, the evolved cells reach exponential-phase growth faster than the ancestor, owing to changes in growth rates and lag times (41). Also, cell densities are lower during exponential growth, especially given the low glucose concentration in the LTEE medium. Nonetheless, we performed a set of experiments to compare the average volumes of exponentially growing and stationary-phase populations (Fig. 5). Exponentially growing cells were larger than stationary-phase cells, and this difference was observed using both the ancestor and evolved bacteria, Bacterial cells are larger during exponential growth to accommodate more ribosomes (18) and replicating chromosomes (13, 17). The approximately two-fold difference in average cell volume between exponential and stationary phases for the 50,000-generation clones (Fig. S2) implies that these bacteria undergo a reductive division as they enter or during stationary phase.

The 12 LTEE populations have evolved shorter lag phases and faster maximal growth rates during their adaptation to the LTEE environment. Therefore, when compared to the ancestor, evolved cells spend more time in the stationary-phase period between transfers. In silico models of the daily transfer regime typical of experimental evolution systems, including the LTEE, have shown that virtual microbes can evolve to anticipate the transfer interval (62). A reductive division during stationary phase might prime the cells to grow faster when transferred into fresh medium by temporarily increasing their SA/V ratio, potentially reducing the duration of the lag phase. However, we note that the LTEE ancestors also undergo a similar reductive division, as indicated by smaller cells in stationary phase than during exponential growth (Fig. 5). Thus, the reductive division per se does not account for the shortened lag phase in the evolved bacteria. In any case, future studies might examine when this reductive division occurs in the ancestral and evolved bacteria and, moreover, identify the metabolic cues and physiological processes involved.

We also observed substantial heterogeneity in the cell shape of the evolved lines (Fig. 7). One population (Ara+5) evolved stubby, almost spherical, cells early in the LTEE (Fig. 11A), evidently caused by a mutation in *mreB*, which encodes a protein involved in maintaining the rod shape that is typical of *E. coli*. This population later re-evolved more rod-shaped cells (Fig. 11B), although the genetic basis for that change is unclear. More generally, most populations evolved relatively wider cells during the first 10,000 generations (Fig. 8), even though longer cells would have had a higher SA/V ratio (33). This trend suggests that cell size evolution in the LTEE is not tightly constrained by the SA/V ratio. In later generations, the average cell aspect ratio (length/width) reverted to the ancestral ratio (Fig. 9A), but not enough to prevent a further decline in the average SA/V ratio (Fig. 9B), as the mean cell volume continued to increase (Fig. 3A).

For a given cell volume, wider cells have lower SA/V ratios than longer cells. From the standpoint of acquiring limited nutrients, wider cells would therefore seem maladaptive, yet that is how the LTEE populations tended to evolve for the first 10,000 generations (Fig. 8). Might wider cells have had some benefit that overcame their unfavorable SA/V ratios? As a bacterial cell grows in size, it simultaneously replicates multiple copies of its chromosome. These copies must then be fully segregated into the two daughter cells, which requires moving them away from the cell center before the division can be completed (13). Rod-shaped bacteria like *E. coli* typically divide at the middle of the cell; the midpoint is defined by the proteins MinCDE, which oscillate between the cell poles every 40-90 seconds while consuming ATP (63, 64, 65, 66, 67, 68). The number of MinCDE complexes doubles in cells longer than ∼4 µm, while their oscillatory period remains constant (63). It has also been shown that MinCDE proteins do not oscillate at all in shorter cells, which have a reduced aspect ratio; instead, they exhibit stochastic switching between the two poles (69). This stochastic switching reduces the rate at which these proteins use ATP (70). Thus, one could imagine that evolving wider cells, which also have a reduced aspect ratio, would increase the ATP available for other metabolic processes. Future studies might study the oscillatory behavior and ATP consumption of these proteins in the LTEE lines.

Another potential advantage of wider cells is to minimize the macromolecular crowding that occurs within the highly concentrated cellular cytoplasm (71). Gallet et al. (72) suggested that the increased cell width in the LTEE lines might reduce the adverse effects of macromolecular crowding, but they did not directly test this hypothesis. However, they also proposed that the bacterial cells became larger in order to become less densely packed, which would allow greater internal diffusion of resources and macromolecules. Gallet et al. (72) found evidence in support of this second hypothesis in the one LTEE population they examined, where the cell density (dry mass-to-volume) declined over evolutionary time. If the rate of resource acquisition from the external environment does not limit growth, then increasing the rate of internal diffusion should increase the cell’s metabolic rate and, at least potentially, lead to faster growth and higher fitness (1, 73, 74, 75, 76, 77, 78, 79). Therefore, it would be interesting to extend the analyses performed by Gallet et al. (72) to all of the LTEE populations to assess the generality of their findings.

We also made the serendipitous discovery that one population, called Ara−3, evolved greatly elevated cell mortality (Figs. 10 and 19). That population is the only one that evolved the ability to assimilate energy from citrate, which is in the LTEE medium as an iron chelator (47). We subsequently showed that this increased mortality has persisted in the population for almost 20,000 generations, and perhaps even longer (49). The persistence of this elevated death suggests some physiological constraint that is difficult to overcome, though this cost must be smaller than the benefit provided by the access to this additional resource. In any case, a 50,000-generation clone that we analyzed from this population was also an outlier in other morphological respects, producing cells that are exceptionally large (Fig. 2A) and long (Fig. 8). In addition to the many ghost-like cells that appear to be dead or dying (Fig. 7), some of these translucent cells have inclusions within the cytoplasm (Fig. 14). Future studies may investigate the genetic and physiological bases of these unusual morphological traits and their relation to growth on citrate and cell death.

In summary, we have observed substantial changes in cell morphology, including shape as well as size, over the course of 50,000 generations of the *E. coli* LTEE. Some of the changes are highly repeatable including especially the parallel trend toward larger cells observed in all 12 independently evolving populations. At the same time, the replicate populations have evolved highly variable phenotypes, even under identical conditions, leading to approximately two-fold variation in their average cell volumes (Fig. 5) as well as large differences in their aspect ratios (Fig. 8). The consistent trend toward larger cells (Fig. 2B), the strong positive correlation of cell volume with fitness (Fig. 13), and the parallel substitutions in genes involved in maintaining cell shape (Fig. 12) all suggest that the evolution of cell morphology is not a mere spandrel, but instead reflects adaptation to the LTEE environment. The resulting among-population variation in size and shape, however, suggest that precise changes in cell morphology were not critical to performance, because most populations have improved in relative fitness to a similar degree (43), despite different cell morphologies. Thus, the changes in cell size and shape during the LTEE reflect both natural selection and the idiosyncratic nature of the chance events, including mutations, particular to every evolving lineage.

## Materials and Methods

### Strains

The *E. coli* LTEE is described in detail elsewhere (39, 42, 44). In short, 12 populations were derived from a common ancestral strain, REL606. Six populations descend directly from REL606. The other six descend from REL607, which differs from REL606 by two selectively neutral mutations (55). Whole-population samples and clones from each population have been frozen at 500-generation intervals. These materials permit the retrospective analysis of genotypic and phenotypic evolution. In this study, we used both clones (Table S1) and whole-population samples (Table S2) from 2,000, 10,000 and 50,000 generations.

### Culture conditions

Samples from the freezer were slightly thawed, inoculated into LB broth, and grown overnight at 37°C. These cultures were diluted 1:10,000 in 9.9 mL Davis Mingioli medium containing 25 ug mL^-1^ glucose (DM25). Cultures were incubated at 37°C in 50-mL Erlenmeyer flasks, with orbital shaking at 120 rpm for 24 h. These conditions are the same as those used in LTEE. The following day, we diluted cultures 1:100 in fresh DM25 and grew them for 2 h or 24 h for exponential and stationary phase cell measurements, respectively.

### Volumetric and shape measurements

Cell sizes were measured using two analytical approaches. In one, we used the Coulter Multisizer 4e (Beckman), an electronic device that measures cell volume following the Coulter principle (80). In this study, we used a 30-µm aperture, and we measured particle sizes in the range from 2% to 60% of the aperture diameter, which corresponds to a volumetric range of 0.113 fL to 3,054 fL. However, we excluded any particles over 6 fL in our analyses. On several occasions we calibrated the aperture using 5.037-µm diameter wide latex beads (Beckman). The measured variance in bead size was below the recommended threshold of 2.0% at each calibration.

In the second approach, we imaged cells using phase-contrast microscopy, and we processed the resulting micrographs using the *SuperSegger* package (52). *SuperSegger* automatically identifies the boundary between cells and segments the individual cells on a micrograph. It returns measurements aligned to the midline of each cell for the long and short axes, which we used as length and width, respectively. The volume (in arbitrary units) of a cell is approximated by integrating over all segments within the cell’s boundaries (82). Given the low density of cells in DM25 even at stationary phase, and to obtain sufficient numbers of cells for analysis in many visual fields, we concentrated most cultures 2-fold by centrifugation at 7,745 g for 2 min. Clones from two populations at generation 50,000 (Ara−1 and Ara−4) required 4-fold concentration. Samples from another population at generation 50,000 (Ara−3) were imaged without concentrating the medium. We then spotted 3-µl samples from each processed culture onto 1% agarose pads, and we imaged the cells using a Nikon Eclipse Ti-U inverted microscope.

### Analysis of cell mortality in population Ara−3

We reanalyzed data on cell viability collected for two clones: the LTEE ancestor (REL606), and the 50,000-generation clone from population Ara−3 (REL11364) that evolved the novel ability to use citrate as a source of carbon and energy (Cit^+^). We used the *Bac*Light viability kit for microscopy (ThermoFisher #L7007) following the manufacturer’s directions for fluorescently labeling cells. In short, we mixed the provided components A and B in equal amounts, added 1 µl to 10-mL stationary-phase DM25 cultures of each clone, and incubated them for 20 min in the dark to prevent photobleaching. The two components contain two fluorescent dyes that differentially stain presumptively live and dead cells. For the Cit^+^ clone only, we also examined cells in DM0 medium, which contains the same concentration of citate as DM25, but no glucose. Full methods and additional results in the context of other work are reported in Blount et al. (49).

### Genomic and fitness data

We integrated our analyses of cell size and shape with previously published datasets on the fitness of the evolved bacteria relative to their ancestor, and on the mutations present in the various clones obtained by sequencing and comparing the evolved and ancestral genomes. The fitness data were previously collected by Wiser et al. (42), who performed competition assays between evolved populations and reciprocally marked ancestors. We downloaded these data from the Dryad Digital Repository (accession https://doi.org/10.5061/dryad.0hc2m). The complete genomes of the ancestral strain and evolved clones used in our study were sequenced by Jeong et al. (81) and Tenaillon et al. (55), respectively. We used an online tool (http://barricklab.org/shiny/LTEE-Ecoli/) to identify all of the mutations that occurred in several genes (*mreB, mreC, mreD, mrdA,* and *mrdB*) known to be involved in maintaining rod-shaped cells in *E. coli*.

### Statistical analyses

Statistical analyses were performed in R (Version 3.5.0, 2018-04-23). Our datasets and R analysis scripts will be made available on the Dryad Digital Repository (DOI pending publication).

## Acknowledgments

We thank Terence Marsh, Charles Ofria, Gemma Reguera, and Chris Waters for feedback as this research progressed, and members of the Lenski lab for valuable discussions. We also thank Rohan Maddemsetti and Zachary Blount for their comments on the manuscript. This work was supported in part by a grant from the National Science Foundation (currently DEB-1951307), the BEACON Center for the Study of Evolution in Action (DBI-0939454), and the USDA National Institute of Food and Agriculture (MICL02253). Any opinions, findings, and conclusions or recommendations expressed in this material are those of the authors and do not necessarily reflect the views of the funders.

## References

1. Marshall WF, Young KD, Swaffer M, Wood E, Nurse P, Kimura A, Frankel J, Wallingford J, Walbot V, Qu X, Roeder AHK. 2012. What determines cell size? BMC Biol 10:101.

2. Young KD. 2006. The selective value of bacterial shape. Microbiol Mol Biol Rev 70:660–670.

3. Corno G, Jürgens K. 2006. Direct and indirect effects of protist predation on population size structure of a bacterial strain with high phenotypic plasticity. Appl Environ Microbiol 72:78–86.

4. Batani G, Pérez G, Martínez de la Escalera G, Piccini C, Fazi S. 2016. Competition and protist predation are important regulators of riverine bacterial community composition and size distribution. J Freshw Ecol 31:609–623.

5. Champion JA, Walker A, Mitragotri S. 2008. Role of particle size in phagocytosis of polymeric microspheres. Pharm Res 25:1815–1821.

6. Doshi N, Mitragotri S. 2010. Macrophages recognize size and shape of their targets. PLoS One 5:e10051.

7. St-Pierre F, Endy D. 2008. Determination of cell fate selection during phage lambda infection. Proc Natl Acad Sci U S A 105:20705–20710.

8. Choi C, Kuatsjah E, Wu E, Yuan S. 2010. The effect of cell size on the burst size of T4 bacteriophage infections of *Escherichia coli* B23. JEMI 14:85–91.

9. Koch AL. 2003. Bacterial wall as target for attack: past, present, and future research. Clin Microbiol Rev 16:673–687.

10. Miller C. 2004. SOS response induction by beta-lactams and bacterial defense against antibiotic lethality. Science 305:1629–1631.

11. Nikolaidis I, Favini-Stabile S, Dessen A. 2014. Resistance to antibiotics targeted to the bacterial cell wall. Protein Sci 23:243–259.

12. Cooper GM. 2000. The Cell: A Molecular Approach, 2nd edition. Sinauer, Sunderland, Mass.

13. Chien AC, Hill NS, Levin PA. 2012. Cell size control in bacteria. Curr Biol 22:R340–R349.

14. Schaechter M, Maaloe O, Kjeldgaard NO. 1958. Dependency on medium and temperature of cell size and chemical composition during balanced growth of *Salmonella typhimurium*. J Gen Microbiol 19:592–606.

15. Bremer H, Dennis PP. 2008. Modulation of chemical composition and other parameters of the cell by growth rate. Escherichia coli and Salmonella typhimurium: Cellular and Molecular Biology 3:1–2.

16. Akerlund T, Nordstrom K, Bernander R. 1995. Analysis of cell size and DNA content in exponentially growing and stationary-phase batch cultures of *Escherichia coli*. J Bacteriol 177:6791–6797.

17. Wang JD, Levin PA. 2009. Metabolism, cell growth and the bacterial cell cycle. Nat Rev Microbiol 7:822–827.

18. Valgepea K, Adamberg K, Seiman A, Vilu R. 2013. *Escherichia coli* achieves faster growth by increasing catalytic and translation rates of proteins. Mol Biosyst 9:2344–2358.

19. Amir A. 2017. Is cell size a spandrel? ELife 6:1–8.

20. Gould SJ, Lewontin RC. 1979. The spandrels of San Marco and the panglossian paradigm: a critique of the adaptationist programme.Proc R Soc Lond B Biol Sci 205:581–598.

21. Heim NA, Payne JL, Finnegan S, Knope ML, Kowalewski M, Lyons SK, McShea DW, Novack-Gottshall PM, Smith FA, Wang SC. 2017. Hierarchical complexity and the size limits of life. Proc R Soc Lond B Biol Sci 284:20171039.

22. Rappé MS, Connon SA, Vergin KL, Giovannoni SJ. 2002. Cultivation of the ubiquitous SAR11 marine bacterioplankton clade. Nature 418:630–633.

23. Giovannoni SJ. 2017. SAR11 bacteria: the most abundant plankton in the oceans. Ann Rev Mar Sci 9:231–255.

24. Angert ER, Clements KD, Pace NR. 1993. The largest bacterium. Nature 362:239–241.

25. Levin PA, Angert ER. 2015. Small but mighty: cell size and bacteria. Cold Spring Harb Perspect Biol 7:a019216.

26. Mika JT, van den Bogaart G, Veenhoff L, Krasnikov V, Poolman B. 2010. Molecular sieving properties of the cytoplasm of *Escherichia coli* and consequences of osmotic stress. Mol Microbiol 77:200–207.

27. Mika JT, Poolman B. 2011. Macromolecule diffusion and confinement in prokaryotic cells. Curr Opin Biotechnol 22:117–126.

28. Schavemaker PE, Boersma AJ, Poolman B. 2018. How important is protein diffusion in prokaryotes? Front Mol Biosci 5:1–16.

29. Yang DC, Blair KM, Salama NR. 2016. Staying in shape: the impact of cell shape on bacterial survival in diverse environments. Microbiol Mol Biol Rev 80:187–203.

30. Sourjik V, Wingreen NS. 2012. Responding to chemical gradients: bacterial chemotaxis. Curr Opin Cell Biol 24:262–268.

31. Tucker JD, Siebert CA, Escalante M, Adams PG, Olsen JD, Otto C, Stokes DL, Hunter CN. 2010. Membrane invagination in *Rhodobacter sphaeroides* is initiated at curved regions of the cytoplasmic membrane, then forms both budded and fully detached spherical vesicles. Mol Microbiol 76:833–847.

32. Ojkic N, Serbanescu D, Banerjee S. 2019. Surface-to-volume scaling and aspect ratio preservation in rod-shaped bacteria. ELife 8:1–11.

33. Harris LK, Theriot JA. 2018. Surface area to volume ratio: a natural variable for bacterial morphogenesis. Trends Microbiol 26:815–832.

34. Kawecki TJ, Lenski RE, Ebert D, Hollis B, Olivieri I, Whitlock MC. 2012. Experimental evolution. Trends Ecol Evol 27:547–560.

35. Ratcliff WC, Denison RF, Borrello M, Travisano M. 2012. Experimental evolution of multicellularity. Proc Natl Acad Sci U S A 109:1595–1600.

36. Graves JL, Hertweck KL, Phillips MA, Han MV, Cabral LG, Barter TT, Greer LF, Burke MK, Mueller LD, Rose MR. 2017. Genomics of parallel experimental evolution in *Drosophila*. Mol Biol Evol 34:831–842.

37. Lenski RE, Ofria C, Pennock RT, Adami C. 2003. The evolutionary origin of complex features. Nature 423:139–144.

38. LaBar T, Adami C. 2017. Evolution of drift robustness in small populations. Nat Commun 8:1012.

39. Lenski RE, Rose MR, Simpson SC, Tadler SC. 1991. Long-term experimental evolution in *Escherichia coli*. I. Adaptation and divergence during 2,000 generations. Am Nat 138:1315–1341.

40. Lenski RE, Travisano M. 1994. Dynamics of adaptation and diversification: a 10,000-generation experiment with bacterial populations. Proc Natl Acad Sci U S A 91:6808–6814.

41. Vasi F, Travisano M, Lenski RE. 1994. Long-term experimental evolution in *Escherichia coli*. II. Changes in life-history traits during adaptation to a seasonal environment. Am Nat 144:432–456.

42. Wiser MJ, Ribeck N, Lenski RE. 2013. Long-term dynamics of adaptation in asexual populations. Science 342:1364–1367.

43. Lenski RE, Wiser MJ, Ribeck N, Blount ZD, Nahum JR, Morris JJ, Zaman L, Turner C, Wade B, Maddamsetti R, Burmeister AR, Baird EJ, Bundy J, Grant NA, Card KJ, Rowles M, Weatherspoon K, Papoulis SE, Sullivan R, Clark C, Mulka JS, Hajela N. 2015. Sustained fitness gains and variability in fitness trajectories in the long-term evolution experiment with *Escherichia coli*. Proc R Soc Lond B Biol Sci 282:20152292.

44. Lenski RE. 2017. Experimental evolution and the dynamics of adaptation and genome evolution in microbial populations. ISME J 11:2181–2194.

45. Lenski RE, Mongold JA. 2000. Cell size, shape, and fitness in evolving populations of bacteria. In: Brown J, West G (eds), Scaling in Biology. Oxford University Press, Oxford, pp 221–234.

46. Philippe N, Pelosi L, Lenski RE, Schneider D. 2009. Evolution of penicillin-binding protein 2 concentration and cell shape during a long-term experiment with *Escherichia coli*. J Bacteriol 191:909–921.

47. Blount ZD, Borland CZ, Lenski RE. 2008. Historical contingency and the evolution of a key innovation in an experimental population of *Escherichia coli*. Proc Natl Acad Sci U S A 105:7899–7906.

48. Blount ZD, Barrick JE, Davidson CJ, Lenski RE. 2012. Genomic analysis of a key innovation in an experimental *Escherichia coli* population. Nature 489:513–518.

49. Blount ZD, Maddamsetti R, Grant NA, Ahmed ST, Jagdish T, Baxter JA, Sommerfeld BA, Tillman A, Moore J, Slonczewski JL, Barrick JE, Lenski RE. 2020. Genomic and phenotypic evolution of *Escherichia coli* in a novel citrate-only resource environment. ELife 9:e55414.

50. Mongold JA, Lenski RE. 1996. Experimental rejection of a nonadaptive explanation for increased cell size in *Escherichia coli*. J Bacteriol 178:5333– 5334.

51. Chang F, Huang KC. 2014. How and why cells grow as rods. BMC Biol 12:54

52. Stylianidou S, Brennan C, Nissen SB, Kuwada NJ, Wiggins PA. 2016. SuperSegger : robust image segmentation, analysis and lineage tracking of bacterial cells. Mol Microbiol 102:690–700.

53. Si F, Li D, Cox SE, Sauls JT, Azizi O, Sou C, Schwartz AB, Ericstad MJ, Jun Y, Li X, Jun S. 2017. Invariance of initiation mass and predictability of cell size in *Escherichia coli*. Curr Biol 27:1278–1287.

54. Kruse T, Bork-Jensen J, Gerdes K. 2005. The morphogenetic MreBCD proteins of *Escherichia coli* form an essential membrane-bound complex. Mol Microbiol 55:78–89.

55. Tenaillon O, Barrick JE, Ribeck N, Deatherage DE, Blanchard JL, Dasgupta A, Wu GC, Wielgoss S, Cruveiller S, Médigue C, Schneider D, Lenski RE. 2016. Tempo and mode of genome evolution in a 50,000-generation experiment. Nature 536:165–170.

56. Pötter M, Steinbüchel A. 2006. Biogenesis and structure of polyhydroxyalkanoate granules. In: Shively JM (ed), Inclusions in Prokaryotes. Springer, Berlin, pp 110– 136.

57. Jendrossek D. 2009. Polyhydroxyalkanoate granules are complex subcellular organelles (carbonosomes). J Bacteriol 191:3195–3202.

58. Al Rowaihi IS, Paillier A, Rasul S, Karan R, Grötzinger SW, Takanabe K, Eppinger J. 2018. Poly(3-hydroxybutyrate) production in an integrated electromicrobial setup: Investigation under stress-inducing conditions. PLoS One 13:e0196079.

59. Obruca S, Sedlacek P, Slaninova E, Fritz I, Daffert C, Meixner K, Sedrlova Z, Koller M. 2020. Novel unexpected functions of PHA granules. Appl Microbiol Biotechnol 104:4795–4810.

60. Rozen DE, Schneider D, Lenski RE. 2005. Long-term experimental evolution in *Escherichia coli*. XIII. Phylogenetic history of a balanced polymorphism. J Mol Evol 61:171–180.

61. Grosskopf T, Consuegra J, Gaffé J, Willison J, Lenski RE, Soyer OS, Schneider D. 2016. Metabolic modelling in a dynamic evolutionary framework predicts adaptive diversification of bacteria in a long-term evolution experiment. BMC Evol Biol 16:163.

62. van Dijk B, Meijer J, Cuypers TD, Hogeweg P. 2019. Trusting the hand that feeds: microbes evolve to anticipate a serial transfer protocol as individuals or collectives. BMC Evol Biol 19:201.

63. Raskin DM, de Boer PA. 1999a. Rapid pole-to-pole oscillation of a protein required for directing division to the middle of *Escherichia coli*. Proc Natl Acad Sci U S A 96:4971–4976.

64. Raskin DM, de Boer PA. 1999b. MinDE-dependent pole-to-pole oscillation of division inhibitor MinC in *Escherichia coli*. J Bacteriol 181:6419–6424.

65. Huang KC, Meir Y, Wingreen NS. 2003. Dynamic structures in *Escherichia coli*: Spontaneous formation of MinE rings and MinD polar zones. Proc Natl Acad Sci U S A 100:12724–12728.

66. Lutkenhaus J. 2007. Assembly dynamics of the bacterial MinCDE system and spatial regulation of the Z ring. Annu Rev Biochem 76:539–562.

67. Dajkovic A, Lan G, Sun SX, Wirtz D, Lutkenhaus J. 2008. MinC spatially controls bacterial cytokinesis by antagonizing the scaffolding function of FtsZ. Curr Biol 18:235–244.

68. Arumugam S, Petrašek Z, Schwille P. 2014. MinCDE exploits the dynamic nature of FtsZ filaments for its spatial regulation. Proc Natl Acad Sci U S A 111:E1192–E1200.

69. Ramm B, Heermann T, Schwille P. 2019. The *E. coli* MinCDE system in the regulation of protein patterns and gradients. Cell Mol Life Sci 76:4245–4273.

70. Fischer-Friedrich E, Meacci G, Lutkenhaus J, Chaté H, Kruse K. 2010. Intra- and intercellular fluctuations in Min-protein dynamics decrease with cell length. Proc Natl Acad Sci U S A 107:6134–6139.

71. Minton AP. 2006. How can biochemical reactions within cells differ from those in test tubes? J Cell Sci 119:2863–2869.

72. Gallet R, Violle C, Fromin N, Jabbour-Zahab R, Enquist BJ, Lenormand R. 2017. The evolution of bacterial cell size: the internal diffusion-constraint hypothesis. ISME J 11:1559–1568.

73. Beveridge TJ. 1988. The bacterial surface: general considerations towards design and function. Can J Microbiol 34:363–372.

74. Koch AL. 1996. What size should a bacterium be? A question of scale. Annu Rev Microbiol 50:317–348.

75. Schulz H, Jorgensen B. 2001. Big bacteria. Annu Rev Microbiol 55:105–137.

76. Golding I, Cox EC. 2006. Physical nature of bacterial cytoplasm. Phys Rev Lett 96:98–102.

77. Beg QK, Vazquez A, Ernst J, de Menezes MA, Bar-Joseph Z, Barabási A-L, Oltvai ZN. 2007. Intracellular crowding defines the mode and sequence of substrate uptake by *Escherichia coli* and constrains its metabolic activity. Proc Natl Acad Sci U S A 104:12663–12668.

78. Ando T, Skolnick J. 2010. Crowding and hydrodynamic interactions likely dominate in vivo macromolecular motion. Proc Natl Acad Sci U S A 107:18457– 18462.

79. Dill KA, Ghosh K, Schmit JD. 2011. Physical limits of cells and proteomes. Proc Natl Acad Sci U S A 108: 17876–17882.

80. Don M. 2003. The Coulter principle: foundation of an industry. JALA 8:72–81.

81. Jeong H, Barbe V, Lee CH, Vallenet D, Yu DS, Choi S-H, Couloux A, Lee S-W, Yoon SH, Cattolico L, Hur C-G, Park H-S, Ségurens B, Kim SC, Oh TK, Lenski RE, Studier FW, Daegelen P, Kim JF. 2009. Genome sequences of *Escherichia coli* B strains REL606 and BL21(DE3). J Mol Biol 394:644–652.

82. Sliusarenko O, Heinritz J, Emonet T, Jacobs-Wagner C. 2011. High-throughput, subpixel precision analysis of bacterial morphogenesis and intracellular spatio-temporal dynamics. Mol Microbiol 80:612–627.

